# Poised cell circuits in human skin are activated in disease

**DOI:** 10.1101/2020.11.05.369363

**Authors:** Gary Reynolds, Peter Vegh, James Fletcher, Elizabeth F.M. Poyner, Emily Stephenson, Issac Goh, Rachel A. Botting, Ni Huang, Bayanne Olabi, Anna Dubois, David Dixon, Kile Green, Daniel Maunder, Justin Engelbert, Mirjana Efremova, Krzysztof Polański, Laura Jardine, Claire Jones, Thomas Ness, Dave Horsfall, Jim McGrath, Christopher Carey, Dorin-Mirel Popescu, Simone Webb, Xiao-nong Wang, Ben Sayer, Jong-Eun Park, Victor A. Negri, Daria Belokhvostova, Magnus Lynch, David McDonald, Andrew Filby, Tzachi Hagai, Kerstin B. Meyer, Akhtar Husain, Jonathan Coxhead, Roser Vento-Tormo, Sam Behjati, Steven Lisgo, Alexandra-Chloé Villani, Jaume Bacardit, Phil Jones, Edel A. O’Toole, Graham S. Ogg, Neil Rajan, Nick J. Reynolds, Sarah A. Teichmann, Fiona Watt, Muzlifah Haniffa

## Abstract

The human skin confers biophysical and immunological protection through a complex cellular network that is established early in development. We profiled ~500,000 single cells using RNA-sequencing from healthy adult and developing skin, and skin from patients with atopic dermatitis and psoriasis. Our findings reveal a predominance of innate lymphoid cells and macrophages in developing skin in contrast to T cells and migratory dendritic cells in adult skin. We demonstrate dual keratinocyte differentiation trajectories and activated cellular circuits comprising vascular endothelial cells mediating immune cell trafficking, disease-specific clonally expanded *IL13/IL22* and *IL17A/F*-expressing lymphocytes, epidermal *IL23*-expressing dendritic cells and inflammatory keratinocytes in disease. Our findings provide key insights into the dynamic cellular landscape of human skin in health and disease.

**One Sentence Summary:** Single cell atlas of human skin reveals cell circuits which are quantitatively and qualitatively reconfigured in inflammatory skin disease.

## Introduction

Human skin undergoes dramatic adaptation as it transitions from a relatively sterile aquatic environment *in utero* to provide mechanical and immunological protection in a non-sterile terrestrial environment. This function requires coordination by specialized cell types that are organized within the unique skin architecture. The vast majority of leukocytes within skin are regionally mobile and also capable of trafficking to and from other parts of the organism through blood and lymphatic vessels (*1*). There is continuous trafficking of leukocytes across the basement membrane (*2*), which separates the epidermis, primarily composed of keratinocytes, from the dermis, comprising stroma, appendageal structures and vessels. Epidermal keratinocytes undergoing differentiation are desquamated over 30-60 days and are replenished by stem cells in the basal layer, whose progeny move upwards to the superficial stratum corneum (*3*).

Recent insights into the origin of immune cells in the skin have suggested seeding of some leukocytes during embryonic development that can persist into adulthood (*4*). The persistence or function of embryonic-derived cells in skin homeostasis or pathology in adult life is unclear.

The contribution of non-leukocytes such as keratinocytes and fibroblasts to cutaneous immunity and immunological memory has revolutionized our understanding of the human skin (*5*). Current consensus view on inflammatory skin disease pathogenesis such as atopic dermatitis (AD) and psoriasis supports the interplay between leukocytes and non-leukocytes in disease initiation and progression (*6*). This knowledge has been primarily derived from whole disease skin transcriptome analysis, which identified the signature pathways for AD (Th2/Th22) and psoriasis (Th17) and their resolution following treatment (*7*). However, the precise cellular interactions mediating these pathways remain unresolved.

Single cell technological advances such as RNA-sequencing provide a new opportunity to dissect the complex cellular organization in human skin in health and disease at a systems level. The limited studies so far have focused on the epidermis (*8*), relatively small cell numbers from dissociated dermis or whole skin (*9*) and have not integrated multiplexed protein analysis. In this study, we used single cell RNA-sequencing (scRNA-seq) to comprehensively define the landscape of the immune networks within early development and healthy adult skin, and characterize the reconfigured cell circuits in two common inflammatory skin disorders, AD and psoriasis.

## Results

### Deconstructing human skin

We performed comprehensive single cell analysis of healthy human skin using scRNA-seq and mass cytometry. To maximize cell yield and viability from tissue dissociation, we used 200μm thick pieces of mammoplasty skin which were separated into epidermis and dermis prior to enzymatic dissociation (**Fig. 1 and Fig. S1A**). We leveraged known surface markers defining the different skin cellular compartments to profile live, single cells from each contiguously placed FACS gates containing non-immune cells (keratinocytes, fibroblasts, and endothelial cells) and immune cells (myeloid and lymphoid cells) (**Fig. S1B**) from the epidermis and dermis for droplet encapsulation (10x Genomics) profiling. We also performed indexed plate-based Smart-seq2 profiling of all epidermal and dermal cells within the CD45^+^HLA-DR^+^ myeloid gate (**Fig. S1B**). To compare cell states in healthy skin with inflammatory disease induced perturbation, we performed scRNA-seq (10x Genomics) on all CD45^+/-^ cells from lesional and non-lesional skin from AD and psoriasis patients (**Fig. 1A and Fig. S1B-E**).

**Figure 1.**
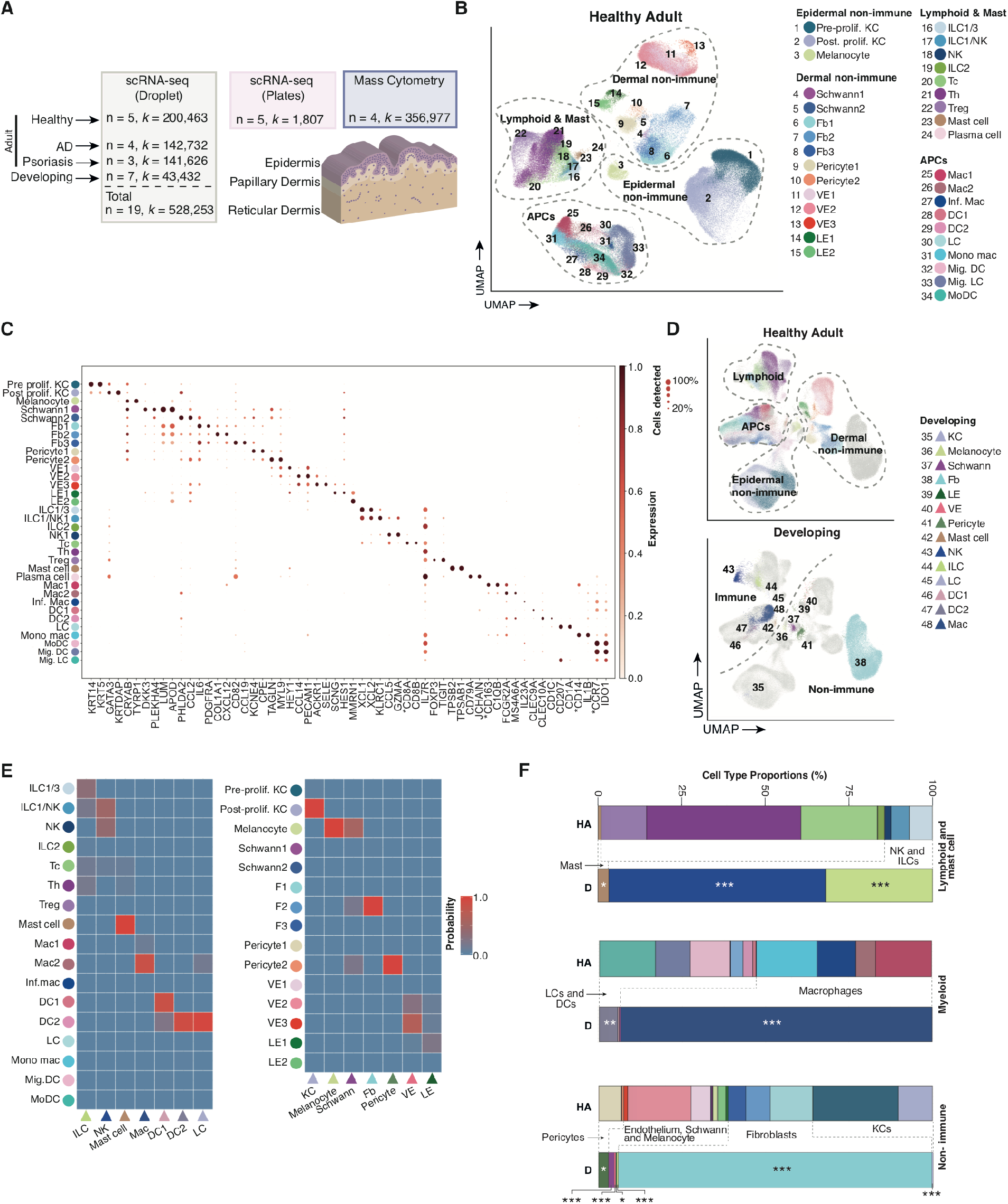
Deconstructing human skin. (a) Number of cells from scRNA-seq and mass cytometry for each condition and schematic of sampling locations. AD = atopic dermatitis (b) UMAP visualization showing all cell states found in the healthy adult scRNA-seq data set, n = 5. Prolif. = proliferative, KC = keratinocyte, Fb = fibroblast, VE = vascular endothelium, LE = lymphatic endothelium, ILC = innate lymphoid cell, NK = natural killer cell, Tc = cytotoxic T cell, Th = T helper cell, Treg = regulatory T cell, Mac = macrophage Inf. = inflammatory, DC = dendritic cell, LC = Langerhans cell, Mono mac = monocyte derived macrophage, Mig. = migratory, MoDC = monocyte derived dendritic cell (c) Dot plot showing the expression of discriminatory markers for each cell state in (b). (d) UMAPs showing the healthy adult cell states found (top) overlaid on the developing cell states (n=7) and the developing cell states (bottom) overlaid on the healthy adult cell states. Cells underlaid are shown in grey. (e) Probability of the cell states found in adult compared to those found in the developing skin using TransferAnchors function in Seurat. (f) Bar charts showing the proportions of each cell states found in adult and developing skin, colors correspond with the legends in (b) and (d). Negative binomial regression comparing developing with adult healthy skin. Statistically significant changes to cell proportion include myeloid cells: DC2, LC and macrophage (*p*<0.001), lymphoid: ILC and NK (*p*<0.001), mast cell (*p*<0.02), fibroblasts, keratinocytes, melanocytes, Schwann cells, VE (*p*<0.001), LE and pericyte (*p*<0.05).

In total, 528,253 sequenced skin cells (*n* = 19) passed quality control (QC) and doublet exclusion (**Fig. 1A and Fig. S2**). We detected on average ~3,000 genes with the 10x Genomics platform and ~6,000 genes per cell with the Smart-seq2 protocol (**see Methods and Fig. S2**). We excluded cells with <200 genes, >20% mitochondrial gene expression and those identified as doublets (**see Methods**). To account for biases due to batch effects we performed data integration of healthy skin samples using BBKNN implementation within Scanpy (*10*, *11*), which showed good mixing by UMAP visualization (**Fig. 1B and Fig. S2A**). We performed graph-based Leiden clustering and derived differentially expressed genes to annotate the cell clusters, from which 34 cell states were identified (**Fig. 1B-C**). We selected genes encoding surface proteins (denoted by * in **Fig. 1C**) and additional antigens where antibodies are commercially available, to derive a CyTOF panel for protein level validation of the cell categories on 4 additional donors (**Fig. S3A-F**). Using CyTOF analysis, we were able to evaluate the frequency of cell types in healthy adult skin (**Fig. S3E**).

To evaluate the establishment of specific cell types during development and the temporal evolution of the cell types found in adult skin, we compared our adult skin scRNA-seq data with our 7-10 post-conception weeks (PCW) (*n* = 7) embryonic/fetal scRNA-seq datasets (*12*). We used the TransferAnchors function in Seurat to integrate adult and fetal skin cell states (**Fig. 1E**) (*13*, *14*) and calculated proportional representation of the equivalent cell states in healthy developing and adult skin (**Fig. 1D-F**).

### Stromal cellular circuits

We next interrogated the heterogeneity within fibroblasts, vascular and lymphatic endothelial cells and Schwann cells (**Fig. 2A-B**). Fibroblasts dominated the non-immune cell population in developing skin with increased proportional representation of melanocytes, Schwann cells and lymphatic and vascular endothelial cells observed in adult skin (**Fig. 1F**).

**Figure 2.**
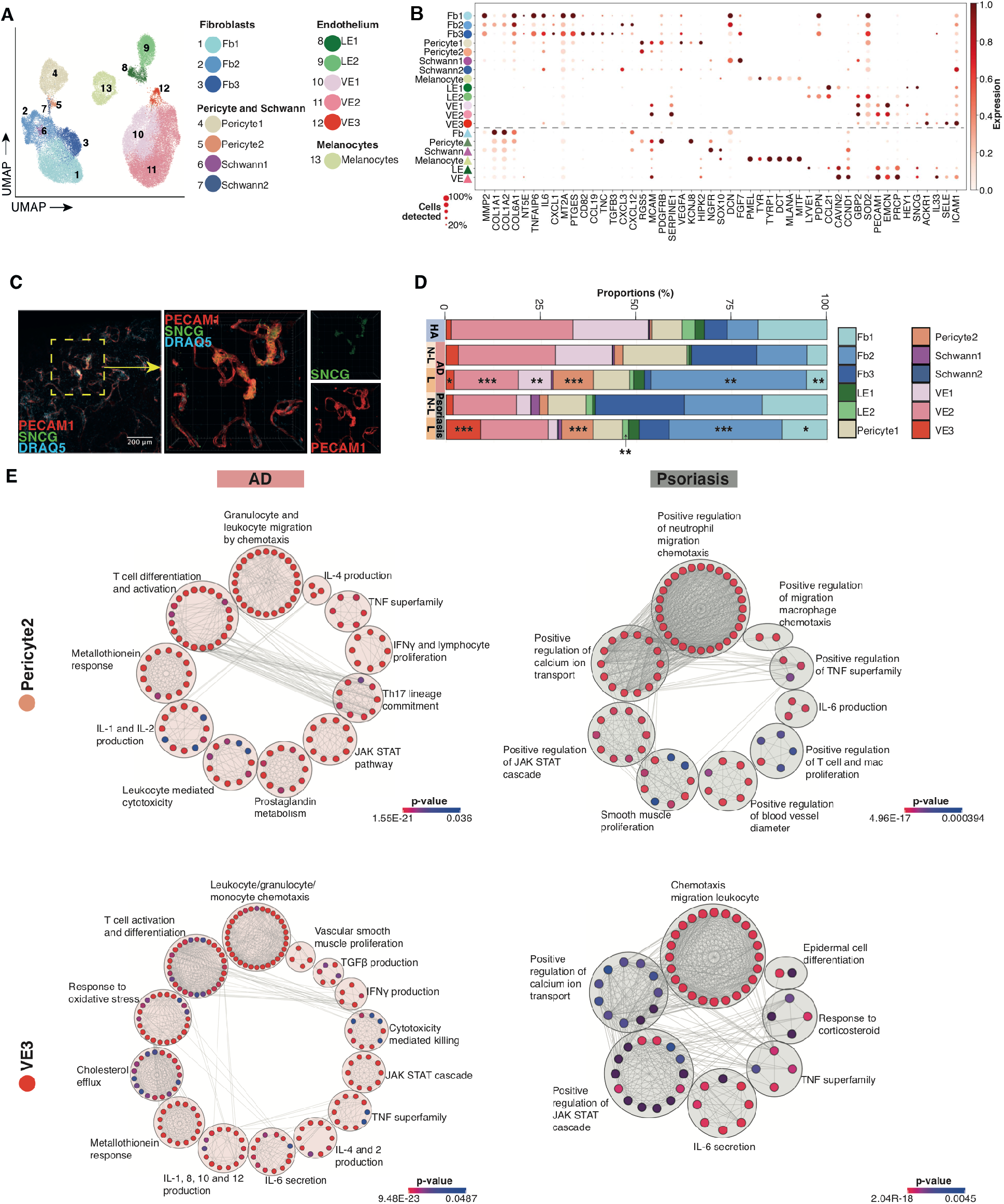
Stromal and endothelial cells. (a) UMAP visualization of the non-immune, non-keratinocyte cell states found in the healthy adult skin, n=5. Fb = fibroblast, LE = lymphatic endothelium, VE = vascular endothelium. (b) Dot plot showing the expression of differentially expressed genes between the cell states found in (a) and their developmental counterparts, separated by the dotted line. (c) 3D Reconstruction of Z-stacked images of whole mount immunofluorescence staining of dermis for CD31 (PECAM1, red), gamma synuclein (SNCG, green) and DRAQ5 (blue). White cubes represent 40×40×40 μm. (d) Bar charts showing the proportions of each cell states found in (a) compared to diseased skin. Negative binomial regression comparing AD and psoriasis with healthy skin. In AD, F1 (*p* = <0.05), F2 (*p* = <0.001), LE2 (*p* = <0.01), Pericyte2 (*p* = <0.001), Schwann1 (*p* = < 0.05), VE1 (*p* = <0.001), VE2 (*p* = <0.01) and VE3 (*p* = <0.001) all significantly change compared to healthy. In psoriasis; F1 (*p* = <0.01), F2 (*p* = <0.01), Pericyte2 (*p* = <0.001), VE1 (*p* = <0.01), VE2 (*p* = <0.001) and VE3 (*p* = <0.05), all significant change compared to healthy. HA = healthy adult, AD = atopic dermatitis, P = psoriasis. (e) Clustered network visualizations of pathways differentially enriched in VE3 and Pericyte2 clusters in AD and psoriasis compared to analogous cell clusters in healthy skin. Network nodes are colored by enrichment score (*p* = <0.05) and represent individual enriched gene sets whilst edges represent shared genes between nodes (intersect >=10%).

Melanocytes are characterized by expression of *PMEL, TYR, TYRP1, DCT1* and *MLANA* (**Fig. 1B-C and 2B**). In fetal skin, these cells express higher levels of *MITF, PMEL* and *TYRP1* but lower levels of *TYR* and *DCT* (**Fig. 2B**) suggesting an immature melanoblast-like profile (*15*). Schwann cells form two clusters, Schwann1 (expressing myelinating genes including *FLG2, NGRF* and *SOX10*) and Schwann2, which co-express Schwann cell markers (*NGRF*) and fibroblast markers (*DCN, FGF7*), and match the DEG profile of cells previously described as a rare fibroblast subset (**Fig. 2B**)(*16*).

Three fibroblast subsets expressing extracellular matrix (ECM)-related genes such as *MMP2, COL1A1, COL1A2* and *NT5E* (encodes CD73) are present in healthy human skin dominated by F1 and F3 fibroblasts and a minor population of F2 fibroblasts (**Fig. 1B-C and 2A-B**). All three also co-express immunomodulatory genes. F1 and F3 fibroblasts express *TNFAIP6*, *IL6* and *CXCL1*, as well as higher levels of *MT2A* and *PTGES* (**Fig. 1C, 2B and Table S1**). F3 fibroblasts share the immunomodulatory gene signature associated with F1 fibroblasts but also express higher levels of

*CD82* (apoptotic), *CCL19* (chemotaxis), as well as wound healing-associated genes including *TNC* and *TGFB3* (*17*) (**Fig. 2B, Table S1**). In contrast to F1 and F3, immunomodulatory genes expressed by F2 fibroblasts include chemokines *CXCL3* and *CXCL12*, involved in monocyte migration and leukocyte inflammation respectively, suggesting additional roles of immune cell recruitment (**Fig. 2B**). Dermal fibroblast heterogeneity encompassing structural and immunomodulatory subtypes has been previously reported at single cell level (*9*, *16*). Interestingly, fibroblasts in fetal skin express more genes relating to F2 fibroblasts, including *COL1A1, COL1A2* and *COL6A1*, suggesting they are functionally specialized towards ECM remodeling and maintenance (**Fig. 2B**). Furthermore, in AD and psoriasis lesional and non-lesional skin, F2 fibroblasts are significantly enriched compared to healthy skin and have upregulated expression of the chemokines *CXCL12* and *CCL19* (**Fig. S4A**).

### Specialized vascular endothelium mediates leukocyte trafficking

Endothelial cells in the healthy adult dermis comprise vascular endothelium (*PECAM1, EMCN*) and lymphatic endothelium (*LYVE1, PDPN*) (**Fig. 2A-B and Table S1**). Pericytes expressing *RGS5, MCAM* and *PDGFRB* and exhibiting mixed endothelium (*SERPINE1*, *VEGFA*) and smooth muscle (*KCNJ8, HIPK2*) features are also present (**Fig. 2A-B and Table S1**).

There are two sub-clusters of lymphatic endothelial cells defined by the differential expression of *LYVE1* and *PDPN*, which are higher in LE1 and LE2 respectively (**Fig. 2A-B**). We previously reported LYVE1 expression to identify initial afferent lymphatics that drain into PDPN^+^ collecting lymphatic vessels in human dermis (*18*). Notably, LE1 cells express higher levels of the chemoattractant *CCL21*, which mediates dendritic cell (DC) migration into skin draining lymph nodes, as well as angiogenesis factors *CAVIN2* and *CCND1*, further supporting their function as initial afferent lymphatics (*19*) (**Fig. 2A-B and Table S1**). LE2 cells can also be differentiated from LE1 by higher expression of *GBP2* and *SOD2* (**Fig. 2B**).

Three distinct states of *PECAM1* (CD31)-expressing vascular endothelial cells are present in adult dermis. *ACKR1* expression in VE2, a marker of venular capillary cells, is distinguishable from *HEY1*-expressing VE1, which correspond to non-venular capillary cells (*20*)(**Fig. 2B**). A third cluster, VE3, which forms ~2% of endothelial cells, is characterized by gamma synuclein (*SNCG*), the venular capillary marker *ACKR1* and inflammatory cytokines, chemokines and leukocyte adhesion molecules including *IL6*, *IL33, SELE* and *ICAM1* (**Fig. 2B**).

We hypothesized that VE3 may demarcate vascular endothelial cells mediating leukocyte adhesion and extravasation into the dermis. We performed whole-mount immunostaining of healthy dermis and identified SNCG^+^PECAM^+^ (VE3) distended vascular structures in the superficial dermis (**Fig. 2C**). Our findings support localized vascular endothelial areas consisting primarily of VE3 specialized in leukocyte adhesion and trafficking (**Fig. 2C**). Notably, *SNCG* expression has been reported in malignant melanoma-associated endothelial cells (*21*), decreased microtubule rigidity (*22*) and cancer metastases (*23*). To further investigate the potential role for VE3 in leukocyte trafficking, we used our receptor-ligand computational framework (CellPhoneDB)(*24*, *25*) to evaluate specifically enriched interactions between VE3 and skin DCs and T cells. VE3 was enriched for the expression of receptors involved in leukocyte adhesion, migration and chemotaxis, in contrast to VE1 and VE2 (**Fig. S4B**).

Both LE and VE are present in 7 PCW fetal skin, earlier than previous studies using 8-10 PCW skin (*26*). Fetal lymphatic endothelial cells have lower expression of *LYVE1, PDPN* and *CCL21* suggesting that they are functionally immature (**Fig. 2B**). Interestingly, fetal VE cells aligned with adult VE3 (**Fig. 1E**) although they do not express *SNCG* (Figure 2B). We observed two pericyte states in healthy adult skin (**Fig. 1B and 2B**). Fetal pericytes aligned with pericyte2 (**Fig. 1E**).

Pericyte2 and VE3 are significantly expanded in AD and psoriasis lesional and non-lesional skin (**Fig. 2D**). To explore the qualitative differences and cell-cell interactions between pericytes and VE3 in healthy and inflamed skin, we interrogated the DEG of pericyte2 and VE3 between AD and psoriasis versus healthy adult skin respectively. Pathway enrichment analysis of these differentially expressed genes revealed shared and distinct gene sets in AD and psoriasis lesional skin pericyte2 and VE3 (**Fig. 2E**). Pericyte2 in AD lesional skin is enriched for leukocyte recruitment and inflammatory responses including TNFα-mediated signaling. Pericyte2 in psoriasis is enriched for neutrophil chemotaxis and stress-response pathways relating to IL-6, and JAK-STAT signaling. VE3 in AD and psoriasis is enriched for leukocyte migration, angiogenesis and stress response pathways. In psoriasis, calcium ion transport-related pathways and lipid biosynthetic responses are also enriched (**Fig. 2E**). As leukocyte chemotaxis featured prominently in the upregulated gene sets in disease, we leveraged CellPhoneDB to establish the receptor-ligand pair interactions between pericyte2 and VE3 and immune cells in AD and psoriasis. which identified *CXCL12:CXCR4* and *SELE/P:SELPLG* involvement (**Fig. S4C**).

### Keratinocyte differentiation in healthy and diseased skin

Further clustering analysis of keratinocytes identified four subgroups corresponding primarily to different proliferative stages of keratinocytes (**Fig. 3A**). Pre-proliferative keratinocytes transcribe basal layer proteins such as *KRT5* and *KRT14* and are abundant in our CD49f^hi^ gate that was used to enrich for basal cells in the epidermis (*27*) (**Fig. 1C**). Our data identifies *CTNNAL1* (alpha-catulin) as a novel marker of pre-proliferative keratinocytes (**Fig. 3B**). In mice, depletion of the related protein, alpha-catenin, upregulated keratinocyte proliferation (*28*). The proliferating keratinocytes have a mixed signature of basal cell transcripts with low level expression of suprabasal cell transcripts, such as *KRT1*. They are characterized by *CDK1* and *PCNA* (**Fig. 3B-C**). Post-proliferative keratinocytes express suprabasal layer genes, such as *KRT1* and *KRT10* (**Fig. 3B-C**). The gene expression patterns of these keratinocyte subgroups are in agreement with their spatial arrangement in the epidermis (**Fig. 3C**)(Human Protein Atlas: www.proteinatlas.org). We observe similar keratinocyte subgroups in a recently published human epidermal cell scRNA-seq dataset (*8*)(**Fig. S5A**). In addition, we identify a cluster of post-proliferative keratinocytes characterized by the expression of immunomodulatory genes, including *CCL20, ICAM1* and *TNF* (**Fig. 3A and C**). In agreement with our analysis of 200μm-deep mammoplasty skin we did not detect hair follicles by histochemical staining (**Fig. S1A**), or bulb and infundibular cells by scRNA-seq, which are found deeper in the skin and are more abundant in other anatomical sites (*29*).

**Figure 3.**
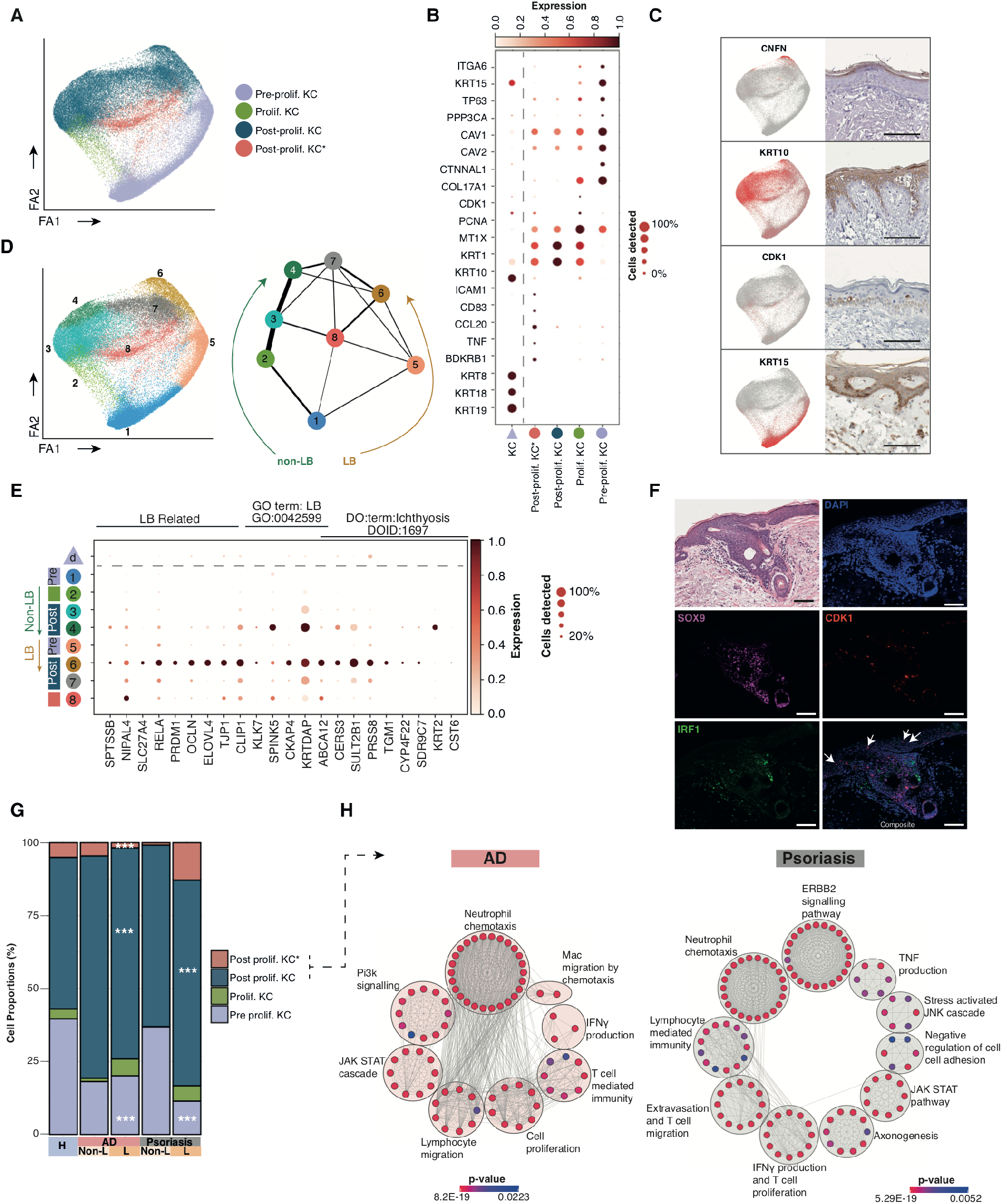
Keratinocyte cell states in health, AD and psoriasis. (a) Force-directed graph (FDG) visualization of the different keratinocyte cell states found in healthy adult skin. Prolif = proliferating, KC = keratinocyte. Asterix indicates the cell state with inflammatory markers, n = 5. (b) FDG feature plots showing gene expression of markers of the cell states found in (a), together with images of these markers *in situ*, from the Human Protein Atlas. Scale bars represent 100μm. (c) Dot plot showing the expression of differentially expressed genes between the cell states found in (a) and their developmental counterparts, separated by the dotted line. (d) Left panel: FDG in (a) annotated by Leiden clustering of eight groups; pre-proliferation (clusters 1, 5), dividing (cluster 2), post-proliferation differentiating (clusters 3, 4, 6, 7, 8). Right panel: PAGA showing the relative connectivity between the keratinocyte clusters. Arrows indicate the two differentiation pathways of basal keratinocytes to suprabasal: LB = lamellar body. (e) Dot plot of genes related to lamellar body production and ichthyosis on the clusters presented in (d), as well as fetal keratinocytes. (f) Immunofluorescence staining of healthy adult skin for DAPI (blue), SOX9 (purple), CDK1 (red), IRF1 (green). Top left image shows an H&E stained sequential section. White arrows indicate CDK1^+^ cells in suprabasal layers Images representative of n = 3. Scale bars represent 100μm. (g) Bar charts showing the proportions of the keratinocyte cell states in healthy and diseased skin. The proportions of post prolif. KC, post prolif. KC* and pre prolif. KC are all significantly different in lesional AD compared to healthy (*p*-values < 0.001). The proportions of post prolif. KC and pre prolif. KC are both significantly different in lesional psoriasis compared to healthy (*p*-values < 0.001). (h) Clustered network visualizations of pathways differentially enriched in post prolif KC clusters in AD and psoriasis compared to analogous cell clusters in healthy skin. Network nodes are colored by enrichment score (*p* = <0.05) and represent individual enriched gene sets whilst edges represent shared genes between nodes (intersect > =10%).

Corresponding analysis of fetal skin shows mixed expression of basal and suprabasal transcripts by fetal keratinocytes (**Fig. 3B**), which comprise two cell layers between 7-10 PCW (**Fig. S5B**). This agrees with histological descriptions of fetal skin comprising an ectodermal derived ‘basal’ layer of undifferentiated keratinocyte progenitors overlaid by the periderm, a transient barrier which is eventually shed into the amniotic fluid (*30*). Fetal keratinocytes express genes for keratins commonly associated with simple epithelium (*31*), including keratin 8, 18 and 19 (**Fig. 3B**). The expression of these keratins declines through gestation and is superseded by the expression of keratins 5 and 14 that characterize adult basal epidermis (**Fig. S5B**).

Force-directed graph (FDG) and partition-based approximate graph abstraction (PAGA) analyses reveal dual inferred differentiation trajectories from the stem cell genes (*TP63, PPP3CA* and *CAV1/2)-enriched* basal cells which transition through proliferating cells into post-proliferative suprabasal cells (**Fig. 3D and Table S1**)(*32*). One arm of keratinocyte differentiation includes cells with low expression of genes associated with the formation of lamellar bodies (LB), which play a key role in late epidermal differentiation (*33*) (**Fig. 3E**). The other arm includes cells with high levels of transcripts involved in LB generation such as *ABCA12, CKAP4* and *CLIP1* (**Fig. 3E**) during suprabasal differentiation. The transcription factors *IRF1* and *SOX9*, differentially expressed in our scRNA-seq analysis by keratinocytes along the two pathways relating to LB-transcript levels (**Fig. S5D**), mark distinct cells by immunofluorescence analysis of healthy skin (**Fig. 3F**). This is consistent with previous findings that human keratinocytes are differentially enriched for lamellar bodies and related proteins (*34*). Transcripts associated with terminal differentiation of keratinocytes (*CNFN, CDSN, FLG* and *IVL*) are enriched in the suprabasal post-proliferative keratinocytes at the end of the differentiation trajectories (**Fig. 3E, Fig. S5C**). The statistically significant differentially expressed genes using Monocle (*35*) across keratinocyte differentiation (**Fig. S5C**) recapitulate some of the previously identified genes in the literature and also in scRNA-seq analysis of murine interfollicular epidermal cells (*36*), including the loss of *KRT5* and *KRT14* and gain of *IVL* and *FLG* from basal to granular layer keratinocytes (*37*). The two pathways are also discernible in AD and psoriasis lesional skin (**Fig. S5E**).

Notably, the suprabasal cells expressing LB-related transcripts in healthy adult skin also coexpress genes associated with autosomal recessive congenital ichthyosis, such as *ABCA12*, *NIPAL4*, *SLC27A4* and *TGM1* (**Fig. 3E**). However, analysis of fetal keratinocytes showed little to no expression of these congenital ichthyosis-related genes, suggesting that disease onset at the molecular level only begins after 10 PCW (**Fig. 3E**). This is in keeping with the absence in fetal epidermis of a granular layer where the expression of *LOR*, *FLG*, *IVL* and genes required for lamellar body production are located (*38*). Notably, AD and psoriasis lesional keratinocytes have altered expression of congenital ichthyosis-related genes due to keratinocyte dysfunction (**Fig. 3E and Fig. S5F**)(*39*).

We observed a cluster of post-proliferative keratinocytes expressing the chemokines *CCL20*, intercellular adhesion molecule *ICAM1, TNF* and the bradykinin receptor *BDKRB1* (**Fig. 3A, C**) in healthy adult but not in developing skin. Strikingly, these cells express higher levels of genes associated with inflammatory ichthyoses and severe atopy such as *NIPAL4* and *SPINK5* (**Fig. 3E**). Both AD and psoriasis lesional skin were enriched for post-proliferative keratinocytes (**Fig. 3G**). The proportion of post-proliferative keratinocytes expressing *CCL20*, *ICAM* and *TNF* are expanded in psoriasis but not AD lesional skin (**Fig. 3B and G**), resembling previously described CCL20-expressing keratinocytes in inflammatory skin diseases induced by Th17 cytokines and TNFα (*40*). Furthermore, lesional AD and psoriasis skin keratinocytes express *KRT6A/B/C* and *KRT16*, psoriasin (*S100A7*), calprotectin (*S100A8/S100A9* heterodimer) serpins (*SERPINB4, SERPINB13*) and the chemotactic factor C10orf99 (**Fig S5G**), previously reported to be expressed in psoriasis and AD (*41*–*43*).

Gene sets enriched in post-proliferative keratinocytes are dominated by immune-mediated pathways in both AD and psoriasis, including lymphocyte migration and TNF and IFNγ signaling (**Fig. 3H**). Notably, the ERBB2 signaling pathway, an epidermal growth factor-like receptor tyrosine kinase, is specifically upregulated in psoriasis (**Fig. 3H**). Improvement in psoriasis has been noted in patients treated with anti-EGFR for underlying carcinoma (*44*).

### Transition from innate to adaptive lymphocytes during skin development

In healthy adult skin both adaptive and innate lymphoid cells are present (**Fig. 4A**). T cells are predominant and consist of the three subtypes, cytotoxic (Tc) (*CD8A/B*), helper (Th) (*CD4*, *CD40LG*) and regulatory (Treg) (*FOXP3, TIGIT, CTLA4*) cells (**Fig. 4A-B**). The majority of these are αβT cells, with a smaller proportion of γδT cells expressing *TRGC2* and *TRDC* found within cytotoxic T cells in the epidermis and dermis (**Fig. S6A**). Innate lymphoid cells (ILCs) were identified by expression of CD161 (*KLRB1*) without co-expression of the delta or gamma subunit of the T cell co-receptor CD3 (*CD3D, CD3G*) (**Fig. 4B**). We identified 4 clusters of innate lymphocytes, consisting of ILC1/3, ILC2, ILC1/Natural Killer (NK) and NK cells (**Fig. 4A**). NK cells expressed *KLRD1, FCGR3A* and the cytotoxic genes *PRF1, GZMB and GNLY*. ILC1/NK have overlapping feature of ILC1 and NKs, as previously described (*45*). Plasticity within ILC1 and ILC3 is also recognized (*46*) as we observed here for skin ILC1/3 (*IL7R, XCL1, XCL2, TNFSRF18 and TNFSRF11*). ILC2 (*IL7R, PTGDR2, GATA3*) has the most homogenous signature in our data and existing literature (**Fig. 4B**) (*47*).

**Figure 4.**
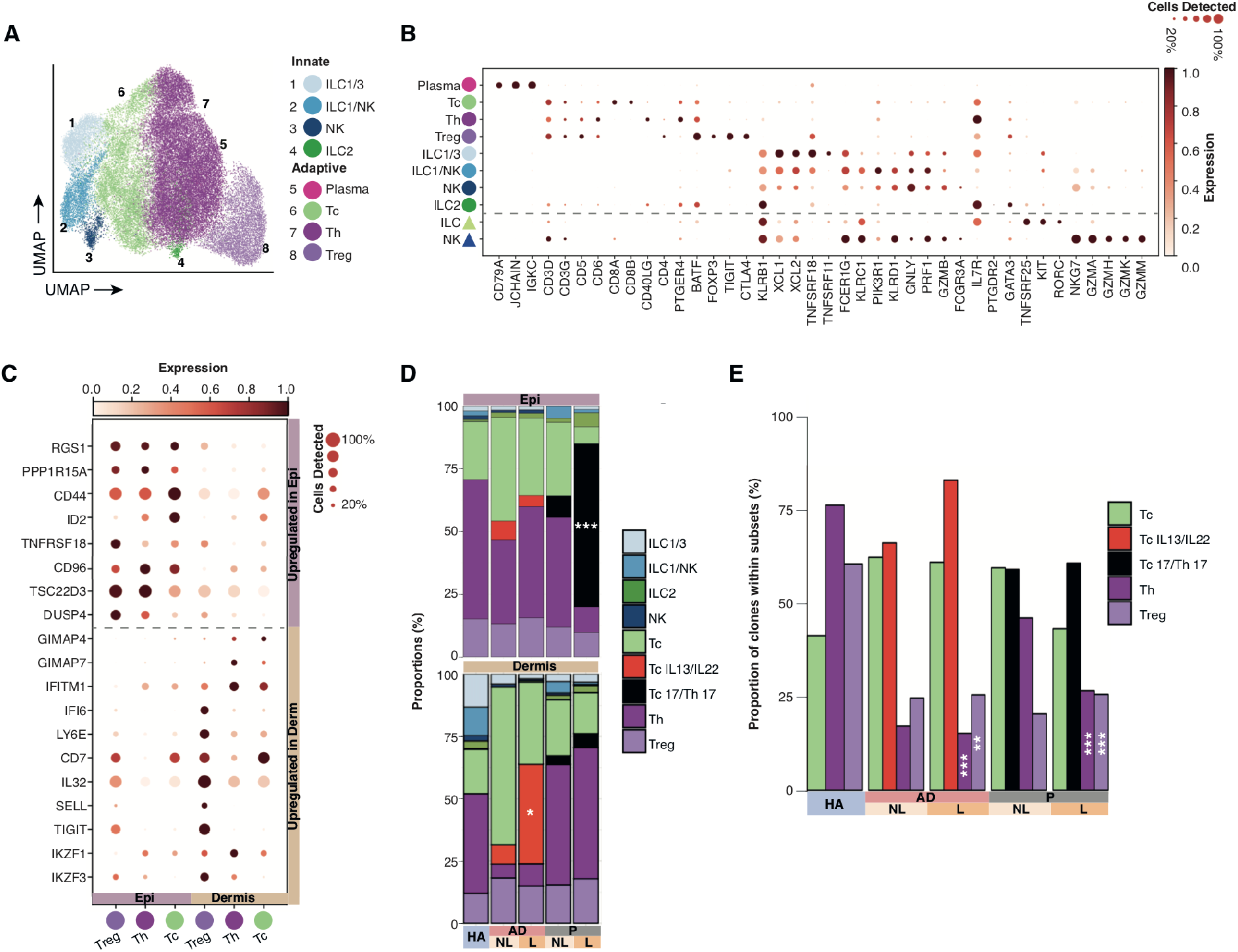
Skin innate lymphoid cells, T lymphocytes and TCR analysis. (a) UMAP visualization of the different lymphoid and mast cell states found in healthy adult skin. ILC = innate lymphoid cell, NK = natural killer cell, Tc = cytotoxic T cell, Th = T helper cell, Treg = regulatory T cell, n = 5 (b) Dot plot showing the expression of differentially expressed genes between the cell states found in (a) and their developmental counterparts, separated by the dotted line. (c) Dot plot showing the differentially expressed genes in the T cell subsets between epidermis and dermis in healthy adult skin, Epi = epidermis, Derm= dermis. (d) Bar charts showing the proportions of the lymphoid cell states in healthy and diseased skin. Lesional dermal AD skin is enriched for TcIL13/IL22 compared to non lesional (*p*-value = 0.04), and psoriasis lesional skin is enriched for Tc17/Th17 in the epidermis (*p*-value < 0.001). Significance of other cell states were not tested. HA = healthy adult, AD = atopic dermatitis, P = psoriasis, L = lesional, NL = non lesional. (e) Bar charts showing the proportion of cells from expanded clonotypes for each T cell subset in health and disease. Proportions are significantly lower for Th (AD *p*-value <0.001, Psoriasis <0.001) and Treg (AD *p*-value <0.01, Psoriasis <0.001) in lesional disease compared to healthy.

In contrast to adult skin, the fetal skin lymphoid compartment is predominantly populated by NK and ILCs (**Fig. 1D**) demonstrating the dominance of innate immune cells prior to the development of the thymus, bone marrow and spleen, where T and B lymphocytes differentiate. Fetal NK cells express *NKG7, GZMA, GZMB, GZMH, PRF1* and low levels of *IL7R* and correlate with adult NK cells (**Fig. 4B**). They also express higher levels of *GZMM* and *GZMK* (**Fig. 4B**), suggesting that they may be functionally competent (*48*). Fetal skin ILCs co-express *IL7R*, *RORC* and *KIT* resembling ILC3 reported in adult skin (**Fig. 4B**) (*49*). Mast cells complete the innate immune compartment of fetal skin and co-express adult mast cell markers such as *KIT*, *CMA1*, *TPSAB1* and *CPA3* (**Fig. 1B-C, Fig. S6B**).

Memory T cells that persist in the skin following an infection acquire properties of resident memory T cells, distinguishing them from central memory T cells found in the peripheral circulation and secondary lymphoid organs (*50*). We compared the gene signatures of skin and blood lymphocytes. Skin lymphocytes aligned with their blood counterparts, but express tissue-residency associated genes such as *CD69* (**Fig. S6C-D**). Dermal T cells align more closely to blood T cells compared to epidermal T cells. (**Fig. S6D**).

To evaluate the impact of the cutaneous compartmental microenvironment on T cells, we compared the DEG between T cells in the epidermis and dermis (**Fig. 4C**). T cells in the epidermis upregulated genes associated with tissue residency (*RGS1, PPP1R15A*)(*57*), effector memory (*CD44, ID2*)(*52*), T cell activation (*TNFRSF18*)(*53*) and inhibition of T cell response (*CD96, DUSP4 and TSC22D3*)(*54*, *55*), in keeping with previous suggestions that resident memory T cells are poised to mount an effective immune response, but express inhibitory molecules to prevent disadvantageous responses to non-pathogenic antigens (*56*). In contrast dermal T cells expressed interferon stimulated genes (*IFITM1, IFI6, LY6E*)(*57*) and genes downregulated in tissue resident T cells (*GIMAP4, GIMAP7*)(*51*). Dermal Treg showed high expression of the central memory T cell marker CD62L (*SELL*), also observed in our CyTOF data (**Fig. S3**), supporting that dermal T cells are more likely to be central memory in contrast to their tissue resident epidermal counterparts.

### Clonal T cells in disease

T cells are known to play a crucial role in the pathogenesis of AD and psoriasis (*58*, *59*). We identified disease-specific T cells expressing cytokine genes associated with AD and psoriasis. In AD, T cells expressing *IL13* and *IL22* (TcIL13/IL22); and in psoriasis, T cells expressing *IL17A* and *IL17F* (Tc17/Th17) are found in both lesional and non-lesional skin (**Fig. 4D and Fig. S6E-F**) but more significantly enriched in lesional skin (**Fig. 4D**). However, they occupy different compartments of the skin, with Tc17/Th17 cells being the dominant T cell subtype in the epidermis of lesional and non-lesional psoriasis (**Fig. 4D**). In contrast, TcIL13/IL22 dominate in the dermis of AD lesional and non-lesional skin (**Fig. 4D**). *IL-13* is expressed in the epidermis of AD, but primarily by ILC2 (**Fig. S6G**), supporting the emerging and integral role of ILC2 in AD (*60*).

The presence and role of epidermal Tc17 cells in psoriasis pathogenesis has been previously reported (*61*, *62*). TcIL13/IL22 cells in AD express amphiregulin (*AREG*), a member of the epidermal growth factor family and genes associated with skin tissue residency (*RGS1, NR4A1, NR4A2*)(*51*), effector T cells (*ID2, PRDM1, MAP3K8*)(*52*, *63*) and T cell activation (*COTL1, MT2A, DUSP2*)(*64*, *65*) (**Fig. S6E**). Tc17/Th17 cells in psoriasis express genes previously reported in activated and pathogenic Th17 cells (*KLRB1, CXCL13* and *RBPJ*)(*66*, *67*), generic T cell activation markers (*JUN, TNFRSF18*)(*53*) and acute stress proteins (*HSPB1, HSPAIB, HSPA1A, DNAJB1, HSPE1*)(**Fig. S6E**).

Significantly more cells belong to expanded Th and Treg clonotypes in healthy skin, compared to psoriasis or AD (**Fig. 4E**). Although the proportion of expanded Tc17/Th17 clonotypes is not significantly different between psoriasis lesional and non-lesional skin, expanded clonotypes are concentrated within the epidermis of lesional skin (**Fig. S6F**).

### Mononuclear phagocytes in adult and developing skin

We next focused on the composition of myeloid cells in adult and fetal skin. We observed 14 states of mononuclear phagocytes (MPs) across both dermal and epidermal layers (**Fig. 5A-B**). Our combined droplet scRNA-seq and Smart-seq2 datasets confirmed heterogeneity of MPs within currently used FACS-gates for these cells (**Fig. S7A-B**). We aligned skin MPs with blood DCs and monocytes using TransferAnchors function in Seurat (**Fig. S7C**) and subset-defining marker genes (*68*) (**Fig. S7D**).

**Figure 5.**
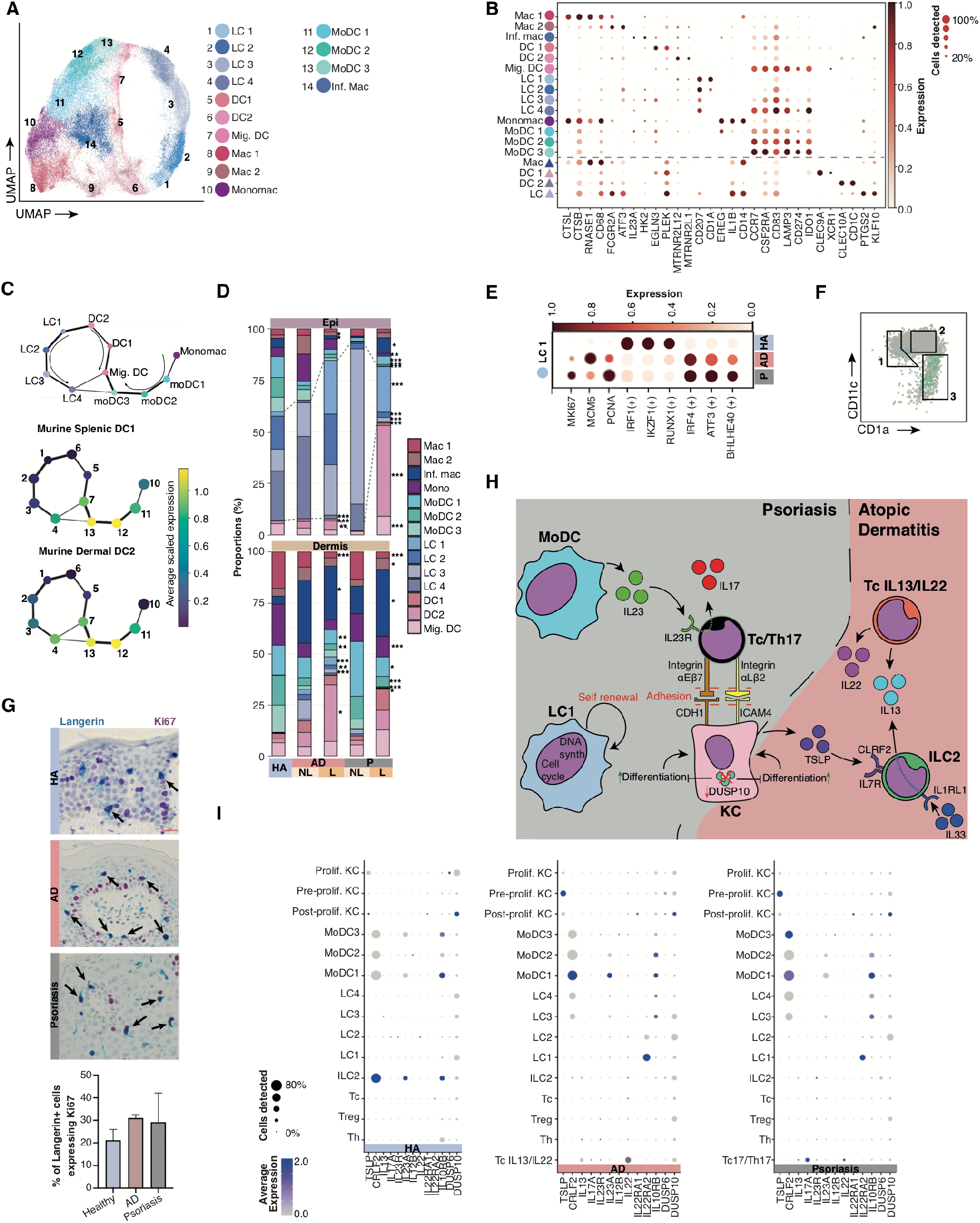
Dermal and epidermal mononuclear phagocytes. (a) UMAP visualization of different antigen presenting cell (APC) states found in adult healthy (n=5), AD (n=4) and psoriasis (n=3). LC = Langerhans cell, DC = dendritic cell, Mig. = migratory, Mac = macrophage, Mono = monocyte derived macrophage, MoDC = monocyte derived dendritic cell, inf. = inflammatory. (b) Dot plot showing the expression of differentially expressed genes between the cell states found (a) and their developmental counterparts, separated by the dotted line. (c) Upper panel: abstracted graph (PAGA) showing connectivity between previously defined clusters. The size of the nodes is proportional to cluster size and edge thickness is proportional to the strength of the connection between nodes. Lower panel: enrichment of gene signatures for murine splenic Xcr1^+^ DC (DC1) and dermal CD11c^+^ (DC2) in each node. (d) Bar charts showing the proportions of MP cell states in healthy, AD and psoriasis skin. In the epidermis proportions of the following cell types significantly change in AD: Mac2 (*p*-value <0.05), Inf. mac (*p*-value <0.05), LC4 (*p*-value <0.001), DC1 (*p*-value <0.001), DC2 (*p*-value <0.01). In the epidermis proportions of the following cell types significantly change in psoriasis: Inf.mac (*p*-value <0.05), mono mac (*p*-value <0.01), moDC1 (*p*-value <0.001), moDC2 (*p*-value <0.001), moDC3 (*p*-value <0.001), LC2 (*p*-value <0.001), LC3 (*p*-value <0.001), LC4 (*p*-value <0.001), DC2 (*p*-value <0.001) and mig. DC (*p*-value <0.001). In the dermis proportions of the following cell types significantly change in AD: Mac1 (*p*-value <0.001), Inf. mac (*p*-value <0.05), moDC1 (*p*-value <0.01), moDC2 (*p*-value <0.01), LC1 (*p*-value <0.001), LC2 (*p*-value <0.01), LC3 (*p*-value <0.001) and DC2 (*p*-value <0.01). In the dermis proportions of the following cell types significantly change in psoriasis: Mac1 (*p*-value <0.001), Mac2 (*p*-value <0.01), Inf. Mac (*p*-value <0.01), monomac (*p*-value <0.001), moDC1 (*p*-value <0.01), moDC2 (*p*-value <0.001), moDC2 (*p*-value <0.001) and LC1 (*p*-value <0.01). (e) Dot plot showing expression of genes associated with proliferation and transcription factors predicted to control the different gene profiles in healthy and diseased skin. (f) Overlay plot of index-sorted epidermal LC1 (light blue) and moDC (green) on all epidermal APCs (live, single, CD45^+^HLA-DR^+^CD3^-^) (grey) (g) Dual stained sections of healthy epidermis stained for Langerin (blue) and Ki67 (purple). Bar chart showing the percentage of Langerin^+^ cells expressing Ki67 in the epidermis of adult healthy skin (n=3), AD (n=4) and psoriasis (n=3). (h) Illustration of predicted cell network interactions active in atopic dermatitis and psoriasis epidermis respectively. (i) Dot plots showing expression of genes forming part of the predicted cellular network in AD and psoriasis (based on (h)).

Two macrophage cell states are present in healthy skin. Mac1 shows higher expression of complement transcripts (*C1QB, C1QC*) and scavenger receptors (*CD163, MARCO*) whereas Mac2 has higher expression of *F13A1* and alternative activation and molecules related to immune suppression (*NR4A1, NR4A2, KLF4*)(**Fig. 5B and Fig. S7D**), and aligned with fetal macrophages (**Fig. 1E**). We also identified *IL23A-expressing* macrophages, which we refer to as inflammatory macrophages (**Fig. 5B**).

We observe DC1, DC2 and Langerhans cells (LCs) in embryonic skin as early as 7 PCW, prior to bone marrow hematopoiesis but with macrophages forming the majority of skin myeloid cells in development (**Fig. 1D-F**). Fetal LCs are enriched for macrophage-related genes such as *CD14, FCGR2A* and *CTSB* (**Fig. 5B**). The embryonic origin of fetal LCs (*69*) and HSC origin of some adult LCs in bone marrow transplant patients (*70*) may explain the poor correlation between fetal and adult LCs (**Fig. 1E**).

### Phenotypic convergence of migratory dendritic cells

In the steady state, skin DCs undergo a continual process of homeostatic maturation that is required to induce tolerance to innocuous environmental antigens (*71*). This is accompanied by their migration to skin draining lymph nodes through lymphatic vessels, a process dependent on CCR7 (*72*). PAGA analysis revealed three branches of differentiation: LCs, myeloid DCs (DC1 and DC2) and monocyte-derived DCs (mo-DCs) (**Fig. 5C**). The clusters at the convergence of these branches (moDC3, LC4 and Mig. DC) express transcripts associated with DC maturation (*CD83, CCR7, LAMP3*) and tolerance (*CD274, IDO1*) (**Fig. 5B and Table S1**). Amongst the top 1000 DEG for each pathway, 33% of genes were common to all three (503 genes) and is enriched for predicted NF-κB targets (105/503 genes predicted to be NF-κB targets, normalized enrichment score 4.225)(*73*) as previously reported in migratory mouse DCs (*71*).

Homeostatic maturation of murine dermal DC2 (*74*) and splenic DC1 (*71*) signature was enriched in LC4, moDC3 and migratory DC (mig. DC) (**Fig. 5C**) suggesting conservation of this signature both across species as well as DC subsets. Furthermore, acquisition of this common gene signature is associated with loss of genes conferring subset identity (**Fig. 5B**), as previously noted (*75*). We observed similar transcripts to those that define our migratory DC (*CCR7*, *IDO1*, *LAMP3*) within migratory DC populations in inflammation in human tonsil, ascites (*76*), lung cancer (*77*) and in our own dataset of rheumatoid arthritis synovial fluid DCs (**Fig. S7E**). This extends the previous observation in mice to human and generalizability of the NF-κB-mediated maturation across species and tissues to all migratory DCs. Interestingly, fetal skin DCs and LCs lack *CCR7* (**Fig. 5B**) and there is no equivalent of the migratory MP populations seen in adult skin (**Fig. 1F**). These observations alongside the immature transcriptional profile of fetal skin lymphatic endothelium suggest that migration to skin draining lymph nodes does not occur prior to 10 PCW.

### APCs in AD and psoriasis skin

In AD there is a significant increase in dermal DC2 (**Fig. 5D**), in the same compartment as TcIL13/IL22 (**Fig. 4D**). DC2 upregulate pathways related to innate immune activation including TLR2, TLR4 and Dectin-1 activation as well as IL-1β signaling (*78*) (**Fig. S8A**). Alongside costimulatory signaling (*CD28:CD80, CD28:CD86, CD40:CD40LG*), the highest predicted interactions between dermal APCs and T cells are multiple co-inhibitory (*CTLA4:CD80*, *CTLA4:CD86*, *PDCD1:CD274*) and anti-inflammatory (*ADORA2A:ENTPD1*, *TGFB1:TGFBR1*) interactions (**Fig. S8B**), consistent with a pathogenic role of these interactions in checkpoint inhibitor induced spongiotic dermatitis (*79*). Inflammatory macrophages are also relatively increased in both AD and psoriasis (**Fig. 5D**).

Epidermal DC2 are enriched in both psoriasis and AD skin. Highly enriched pathways are related to interferon gamma signaling and antigen presentation but also include pathways related to cellular stress, DNA damage and repair (**Fig. S8C**). Genes differentially expressed by epidermal DCs include the alarmins *HMGB1, HMGN1, ITGB2, S100A8, S100A9*, which are endogenous inflammatory mediators released by damaged cells and previously reported to promote keratinocyte proliferation and cytokine production (*80*) (**Fig. S8D**).

In both AD and psoriasis LC1 have the highest enrichment of cell cycle genes (*81*) (**Fig. 5E and Fig. S8E**). Using FACS index data we show that LC1 are enriched within the Langerin^+^CD1a^lo^CD11c^lo^gate (**Fig. 5F**). In contrast, the epidermal HLA-DR^+^CD1a^-^Lang^-^ CD11c^+^CD1c^+^ cells are predominantly mo-DCs and correspond to the recently described non-LC like epidermal cells potent at stimulating T cell proliferation, pro-inflammatory cytokine production, and transmission of HIV to CD4^+^ T cells (*82*). To validate our findings, we examined LC proliferation in healthy, psoriatic and AD epidermis. Consistent with previous findings (*83*) we found Ki67^+^Langerin^+^ cells increased in AD and psoriasis lesional skin, suggesting the existence of a LC reservoir in healthy tissue with the capacity to perpetuate skin-derived antigen presentation in AD and psoriasis (**Fig. 5G**).

As abnormal keratinocyte activation and differentiation feature in both AD and psoriasis, we integrated our findings of epidermal immune cells and keratinocytes to illustrate the key cellular circuits activated in AD and psoriasis (**Fig. 5H-I**). In psoriasis, epidermal Tc17/Th17, DC2, LC1 and mo-DC expressing *IL23*, appear to mediate critical interactions that culminate in keratinocyte dysfunction (**Fig. 5H-I, Fig. S8F**). In AD these interactions are orchestrated by *IL13*-expressing ILC2 and TcIL13/IL22 cells (**Fig. 5H-I, Fig. S8F**). DUSP10, previously shown to inhibit keratinocyte differentiation (*32*), is significantly downregulated in both AD and psoriasis, contributing to epidermal hyperplasia (**Fig. 5H-I**).

## Discussion

In this study we deployed scRNA-seq to resolve the cellular heterogeneity and organization of healthy human developing skin, adult skin and lesional and non-lesional skin of patients with AD and psoriasis. We reveal the dominance of innate lymphocytes during development, in contrast to T cells in adulthood and the presence of DCs and LCs as early as 7 PCW of life. We identify specialized endothelial cells that express leukocyte adhesion molecules and form dilated capillary vasculature for leukocyte trafficking and pericytes that contribute to AD and psoriasis pathogenesis, and keratinocyte subsets and differentiation trajectories in inflamed skin. We also identified disease-specific clonally expanded TcIL13/IL22 and Tc17/Th17 cells in AD and psoriasis skin respectively with shared TCR expression across the epidermis and dermis of lesional and non-lesional skin. We demonstrate the transcriptomic convergence of DCs and LCs into a migratory DC molecular phenotype reflecting homeostatic migration of skin APC into lymphatic vessels. Collectively, our findings demonstrate micro-anatomically organized immunological niches within healthy human skin which become reconfigured in AD and psoriasis.

Two models for keratinocyte differentiation have been proposed: random division of a single population of stem cells (stochastic) or hierarchical differentiation of stem cells or stem cells and transit-amplifying cells prior to differentiation (hierarchical)(*84*). Our analysis shows that keratinocyte differentiation trajectory involves two pathways present in healthy and inflamed skin. This is consistent with keratinocyte differentiation according to both the stochastic and hierarchical models (*85*, *86*). It is notable that *IRF2*, which is expressed at a higher level in the second pathway involving lamellar body keratinocytes, was reported as a critical regulator of keratinocyte lamellar body proteins in mice (*87*). Although in fetal skin a periderm layer provides protection in an aquatic environment (*88*), lamellar body-related genes are not prominent and post-proliferative inflammatory keratinocytes are absent. The post-proliferative inflammatory keratinocytes expanded in psoriasis lesional skin likely correspond to previously described CCL20-secreting keratinocytes following inflammatory perturbation (*40*) and may be relevant to the Koebner phenomenon in psoriasis.

The specialized vascular endothelial cells (VE3) that form distended capillaries in the dermis morphologically resemble high endothelial venules (HEVs) in lymphoid organs. HEVs mediate leukocyte entry into lymph nodes for immune surveillance and express many of the same leukocyte adhesion and chemokine molecules as VE3 such as *CD34*, *ICAM1* and *IL33* (*89*, *90*). VE3 and pericyte2 in AD and psoriasis appear to be actively involved in leukocyte recruitment and present additional therapeutic targets. Notably, fetal fibroblasts, pericytes and VE cells are aligned to adult counterparts (Fb2, Pericyte2 and VE3) that are relatively expanded in both AD and psoriasis. This suggests that tissue modelling during development may be co-opted in tissue re-modeling in disease.

The striking presence of innate lymphoid cells such as NK cells, DCs and LCs as early as 7 PCW in a sterile *in utero* environment raises the question of their *raison d’être* during early development when afferent lymphatics are poorly developed and T cells are absent. NKs and DCs during early development may play a critical role in tissue genesis and modelling, potentially through interaction with other skin cells via HLA-E, which is known to inhibit killing by NK cells under some circumstances (*91*). The precise molecular dissection of how NK and DCs contribute to tissue development merits further investigation in future studies. Differences in the embryonic or fetal and adult microbiomes in human tissues have been previously reported and may influence the immune composition of the skin as a barrier tissue (*92*, *93*). We identify for the first time that mo-DCs are abundant in healthy human skin. Future work is required to confirm whether skin mo-DCs are blood monocyte- or DC3-derived. The presence of homeostatic maturation and migration of DCs in human skin were previously underestimated as migrating cells downregulate canonical DC and LC markers e.g. CD1a, CD1c and Langerin.

In summary, our comprehensive atlas of human skin reveals key circuits involved in AD and psoriasis pathogenesis and provides a foundational resource on the dynamic cellular topology that evolves during development, adulthood and inflammatory skin disease.

## Acknowledgments

We thank the Newcastle University Flow Cytometry Core Facility, Bioimaging Core Facility, Genomics Core Facility and NUIT for technical assistance, School of Computing for access to the High-Performance Computing Cluster, Newcastle Molecular Pathology Node Proximity Lab and Alison Farnworth for clinical liaison. The human embryonic and fetal material was provided by the Joint MRC / Wellcome (MR/R006237/1) Human Developmental Biology Resource (www.hdbr.org).

## Materials and Methods

RF-10 media consists of Roswell Park Memorial Institute media (RPMI)(Sigma, R0883) supplemented with 10% fetal calf serum (FCS) (Life technologies, 10270106), 100U/ml Penicillin (Sigma, P0781), 100 μg/ml Streptomycin (Sigma, P0781) and 1% (v/v) L-Glutamine (Sigma, G7513). Flow buffer consists of Dulbecco’s phosphate buffered saline (PBS)(Sigma, D8537) supplemented with 2% (v/v) FCS and 2mM EDTA (Sigma, E7889). Wash buffer consists of PBS supplemented with 2% FCS.

### Ethics statement

Patients who donated adult healthy skin and biopsies of atopic dermatitis and psoriasis gave written informed consent, in accordance with the Newcastle and North Tyneside 1 Research Ethics Committee (Newcastle Dermatology Biobank - REC reference: 08/H0906/95+5). Adult healthy skin was donated from normally discarded surplus skin from mammoplasty surgery. Patients with atopic dermatitis and psoriasis consented to a 6mm punch biopsy from lesional and non-lesional skin (at least 2 cm away from lesion) (**Table S2**). All patients were naive to biologics treatment, had been free of any other systemic treatment for at least 4 weeks and free of topical corticosteroids for 1 week. Human embryonic and fetal skin was obtained from the MRC/Wellcome Trust-funded Human Developmental Biology Resource (HDBR; http://www.hdbr.org/)(*94*) with appropriate written consent and approval from the Newcastle and North Tyneside NHS Health Authority Joint Ethics Committee (08/H0906/21+5). HDBR is regulated by the UK Human Tissue Authority (HTA; www.hta.gov.uk) and operates in accordance with the relevant HTA Codes of Practice.

### Fetal developmental stage assignment and chromosomal assessment

Embryos up to 8 post conception weeks (PCW) were staged using the Carnegie staging method (*95*). After 8 PCW, developmental age was estimated from measurements of foot length and heel to knee length and compared against a standard growth chart (*96*). A piece of skin, or where this was not possible, chorionic villi tissue was collected from every sample for Quantitative Fluorescence-Polymerase Chain Reaction analysis using markers for the sex chromosomes and the following autosomes 13, 15, 16, 18, 21, 22, which are the most commonly seen chromosomal abnormalities. All samples were karyotypically normal.

### Generation of single cell suspension from adult skin

Healthy skin was cut to thin strips in PBS and the top 200 μm layer was taken using a dermatome with a Pilling Wecprep blade and a .008 gauge Goulian guard. A grid of slits was cut into the skin sheets to aid enzymatic access and the skin was then treated with 2U/ml dispase II (Roche, 04942078001) in RPMI at 37°C for 1 hour. The epidermis was peeled from the dermis, and both fragments were separately digested in a petri-dish at 37°C 5% CO2 in RF-10 media with 1.6 mg/ml type IV collagenase (Worthington, CLS-4) overnight (12 hours). All work was done within class II biological safety cabinets using autoclave-sterilized equipment. The media was then collected with a serological pipette and filtered through a sterile 100 μm cell strainer (BD Falcon, 352360). The petri dish and strainer were washed through with RF-10 to collect any remaining cells. Cells were pelleted by centrifuging at 500 x g for 5 minutes. Supernatant was discarded and the pellet resuspended in 1 ml RF-10 by gently pipetting up and down. Cells were then counted by taking off 10 μl, mixing 1:1 with 0.4% trypan blue (Sigma, T8154) to stain dead cells and counting on a hemocytometer. The 6 mm punch biopsies from AD and psoriasis patients’ skin were trimmed of the lower dermis and subcutis, and cut in half using a scalpel. The biopsies were then treated with 2 μl of dispase II in RPMI at 37°C for 2 hours. The epidermis was peeled from the dermis, and the remaining tissue processing was the same as written above for healthy skin.

### Flow cytometry analysis of cell suspensions

Suspensions of dermal or epidermal cells were resuspended in 100 μl of flow buffer per 10^7^ cells, transferred to polystyrene FACS tubes (BD Falcon, 352054) and stained with 5 μl of each antibody (**Table S3**) per 10^7^ cells for 30 minutes at 4°C in the dark. Cells were washed by diluting in flow buffer and centrifuging at 500 x g for 5 minutes, then resuspended in 200 μl of flow buffer per 10^6^ cells with 3 μM DAPI (Sysmex Partec, 05-5005). Cells were then run through a Fortessa X20 for analysis. Flow cytometry analysis was performed with FlowJo V10 (FlowJo LLC, USA), and the plots were prepared with GraphPad Prism 7.00 (GraphPad Software, La Jolla California USA).

### Single cell sorting of skin cells

Suspensions of dermal or epidermal cells were resuspended in 100 μl of flow buffer per 10^7^ cells, transferred to polypropylene FACS tubes (BD Falcon, 352063) and stained with 5 μl of each antibody per 10^7^ cells for 30 minutes at 4°C in the dark. Cells were washed by diluting in flow buffer and centrifuging at 500 x g for 5 minutes, then resuspended in 1 ml of sort buffer per 20×10^6^ cells with 3 μM DAPI. Cells were then filtered using a 100 μm cell strainer and sorted using a FACSFusion Sorter with a 100 μm fluidics nozzle.

### Mass cytometry analysis of cell suspensions

Suspensions of dermal or epidermal cells were incubated with 2.5 μM Cell-ID Cisplatin (Fluidigm, 201064) for 5 min at RT before washing twice in wash buffer at 500 x g for 5 minutes. Cell-surface antigens were labelled with a master mix of metal-tagged antibodies (**Table S3**) for 1 hour at room temperature in wash buffer. After washing twice in PBS (500 x g, 5 minutes), cells were fixed for 30 minutes at RT in a solution of 0.5x Fix Buffer (OXP3 buffer set, eBiosciences, 00-5523-00) supplemented with 1.6% formaldehyde (Polysciences, 18814). Post-fixation, the cells were washed twice in permeabilization buffer (FOXP3 buffer set, eBiosciences, 00-5523-00) and subsequently incubated with an intracellularly targeted antibody cocktail for 1 hour at RT. Unbound antibody was removed by washing twice in PBS prior to incubating the cells in Iridium intercalator (Fluidigm, 201192A) at 125 nM in PBS for 1 hour. A final fix was performed in 1.6% formaldehyde for 30 minutes at RT, washed twice in PBS and twice in dH2O. Cell pellets were finally resuspended at 0.5 x10^−6^ cells/mL in dH2O with 10% (v/v) EQ beads (Fluidigm, 201078) for analysis on the Helios mass cytometer (Fluidigm).

### Single cell RNA-seq

For the droplet-encapsulation scRNA-seq experiments, 7000 live, single cells were loaded onto each channel of the Chromium chip (10x Genomics, Pleasanton, CA, USA) before droplet encapsulation on the Chromium Controller. Sequencing libraries were generated using the Single Cell 3’ or 5’ plus TCR enrichment reagent kits as per the manufacturer’s protocol. Libraries were sequenced using an Illumina HiSeq 4000 using v4 SBS chemistry to achieve a minimum depth of 50,000 raw reads per cell using the following parameters: Read1: 26 cycles, i7: 8 cycles, i5: 0 cycles; Read2: 98 cycles.

For the plate-based scRNA-seq experiments, a slightly modified Smart-seq2 protocol was used as previously described (*68*). After cDNA amplification, libraries were prepared (384 cells per library) using the Illumina Nextera XT kit (Illumina Inc, San Diego, CA, USA). Index v2 sets A, B C and D were used per library to barcode each cell before multiplexing. Each library was sequenced to achieve a minimum depth of 1-2 million raw reads per cell using an Illumina HiSeq 4000 using v4 SBS chemistry to generate 75bp paired end reads.

### Whole mount immunofluorescence staining of dermis and epidermis

Peeled dermis and epidermis sheets were cut to 1 cm^2^ and fixed in 2% formaldehyde (Polysciences, 18814) and 30% sucrose in PBS for 24 hours at 4°C. Dermis and epidermis sheets were incubated in 300 mM glycine for 24 hours at 4°C, washed in PBS then blocked and permeabilized in PBS with 20% goat serum (R&D Systems, DY005) and 0.2% Triton X-100 (Sigma, T8787) for 24 hours at 4°C then washed in PBS with 0.1% Triton X-100. Primary antibodies were added in PBS with 20% goat serum and 0.1% Triton X-100 for 48 hours then washed. Secondary antibodies were added in PBS with 20% goat serum and 0.1% Triton X-100 for 24 hours then washed. Spectral DAPI (Perkin Elmer, FP1490) or DRAQ5 (Abcam, ab108410) was added as per manufacturer’s instructions. Sheets were mounted in Vectashield antifade mounting medium (Vector Laboratories, H-1000) and set for at least 24 hours before imaging. Slides were imaged using a Zeiss LSM800 Airyscan/Spinning disk Confocal Microscope and Zeiss Pro software (Zeiss, Germany). Z stack images were reconstructed using Zeiss Pro software.

### Immunohistochemistry

Formalin fixed paraffin embedded blocks of mammoplasty skin were sectioned at 4 μm thickness onto APES-coated slides. Sections were dewaxed for 5 minutes in Xylene (Fisher Chemical) then rehydrated through graded ethanol (99%, 95% and 70%; Fisher Chemical) and washed in running water. Sections were treated with hydrogen peroxide block (1% v/v in water; Sigma) for 10 minutes and rinsed in tap water prior to antigen retrieval, using citrate buffer, pH 6 and pressure heating. Slides were washed with TBS, pH 7.6 for 5 minutes prior to staining. Staining was done using the Vector Immpress Kit (Vector Laboratories). Sections were blot dried and blocked sequentially with 2.5% normal horse serum, avidin (Vector Laboratories) and then biotin (Vector Laboratories) for 10 minutes each and blot dried in between. Sections were incubated for 60 minutes with primary antibody (**Table S3**) diluted in TBS pH 7.6 and washed twice in TBS pH 7.6 for 5 minutes each before incubation for 30 minutes with the secondary antibody supplied with the kit. Slides were washed twice in TBS pH 7.6 for 5 minutes each, and developed using peroxidase chromogen DAB, or an AP substrate for multi-colored staining. Sections were counterstained in Mayer’s Haematoxylin for 30 seconds, washed and put in Scots tap water for 30 seconds. Slides were dehydrated through graded ethanol (70% to 99%) and then placed in Xylene prior to mounting with DPX (Sigma-Aldrich). Sections were imaged on a Nikon Eclipse 80i microscope using NIS-Elements Fv4 or Zeiss LSM800 Airyscan/Spinning disk Confocal Microscope and Zeiss Pro software (Zeiss, Germany). The Human Protein Atlas (version 18.1)(www.proteinatlas.org)(*97*) was used for Figure 3. For a list of URLs pointing to the original photos, see **Table S4**. The Gene Ontology (The Gene Ontology Consortium, 2019)(http://geneontology.org, version: 2019/03/01), the Disease Ontology (*98*) (http://disease-ontology.org, release date: 2019-05-20T12:00:30.342164Z) and the MGD 6.13 (*99*) databases were used to find ichthyosis-related genes.

### Immunohistochemistry Scoring

Dual stained IHC for Ki67 and Langerin were manually counted using a Nikon Eclipse 80i microscope. The number of Langerin^+^ and Ki67^+^Langerin^+^ cells were counted in a 3mm section of epidermis for healthy skin, AD and psoriasis. The percentage of dual stained Ki67^+^Langerin^+^ cells out of total Langerin^+^ cells was calculated.

### Data analysis

#### Alignment, quantification and quality control of scRNA-seq data

Droplet-based (10x) sequencing data was quantified using the Cell Ranger Single-Cell Software Suite (version 3.1.0, 10x Genomics Inc) and aligned to the GRCh38 reference genome (official Cell Ranger reference, version 3.0.0). Smart-seq2 sequencing data was aligned with *STAR* (version 2.5.1b), using the STAR index and aligned to the same GRCh38 reference genome. Gene-specific read counts were calculated using *htseq-count* (version 0.10.0). Cells with fewer than 200 detected genes and for which the total mitochondrial gene expression exceeded 20% were removed. Genes that were expressed in fewer than 3 cells were also removed.

#### Doublet detection and exclusion, data normalization, scaling and feature selection

We applied *Scrublet* (v0.2.1), using an exclusion threshold of median plus four times median absolute deviation of the doublet scores (*100*). Data normalization was performed using the *normalize_per_cell* function in scanpy (v1.4.3, and throughout unless stated otherwise) in Python (v3.6.9) to correct for cell-to-cell variation (*11*). Expression values of each gene were scaled and centered using the *scale* function in scanpy. Highly variable genes were detected using the *highly_variable_genes* function in scanpy with minimum cut-off values 0.0125 and 0.5 for expression and dispersion.

#### Data embedding, visualization and clustering

Principal components were calculated using scanpy.pca and adjusted for donor-to-donor variation using the bbknn package (v1.3.2)(*10*) for batch correction. Dimensionality reduction and embedding was performed using Uniform Manifold Approximation and Projection (UMAP) by the scanpy.tl.umap function. The resultant k-nearest neighbors graph was clustered using the Leiden graph-based method (scanpy.tl.leiden)(*101*).

#### Cluster annotation

Differentially expressed genes were calculated using the Wilcoxon sum rank test restricted to genes expressed in at least 30% of cells in either of the two populations compared, and with a fold change cutoff of 0.25 (natural log scale). All *p*-values were adjusted for multiple testing using the Bonferroni correction. Initial annotation was ascribed by comparing these DEGs to published bulk transcription profiles or protein expression of defined cell types. Following this, four major groups (lymphoid cells, myeloid cells, keratinocytes, and other non-immune cells) were subset for further rounds of feature selection, embedding, visualization and clustering using the same pipeline. For a list of top 30 marker genes for each annotated cell type, see **Table S1**.

#### Trajectory analysis

Reconstruction of differentiation pathways of both APCs and keratinocytes was performed using partition-based approximate graph abstraction (PAGA) in Scanpy v1.4 (sc.tl.paga) (Wolf et al., 2019). PAGA represents clusters with nodes, and edge weights (thickness) quantify the connectivity between groups. Connectivity is a measure of the fraction of inter-cluster edges in the full neighborhood graph of single cells. Diffusion pseudotime values were assigned to cells along the resultant differentiation pathways using scanpy (scanpy.tl.dpt). The R (v3.5.2) (R Core Team 2018) Monocle (v2.6.4) *DifferentialGeneTest* function was used to calculate differentially expressed genes (adjusted *p*-value < 0.001) along each differentiation pathway as a function of their progression through pseudotime (*35*). Transcription factors were identified using The Human Transcription Factor list (v1.01) (*102*).

#### Comparison and correlation between adult and fetal cell types

Adult and fetal datasets were merged in Seurat and both highly variable genes and principal components (PCs) calculated in the combined dataset. To adjust for both sample and stage-related effects the neighborhood graph was calculated with BBKNN using individual samples as the batch key (*10*). UMAP embeddings were calculated based on the BBKNN-derived neighborhood graph and visualized as a scatter plot (**Fig. 1D**). We used the FindTransferAnchors function in Seurat (*13*) to build a correspondence between the fetal (query) and adult (reference) datasets. The resultant prediction scores for each cell were mean averaged by adult clusters and used to produce a heatmap (**Fig. 1E**).

#### V(D)J data analysis

Expanded clonotypes for each cell type were defined as having at least two clones, and the proportions express the ratio of expanded cells to cells with V(D)J data annotation. The Shannon index was calculated to compare clonotype diversity between samples. Each sample was randomly down-sampled to the size of the smallest sample 100 times, and the median of the 100 calculated indices was used. The V(D)J data analysis scripts are available as a Python package (v0.1.2; https://github.com/veghp/pyVDJ).

#### Differences in cell type proportions between conditions

A negative binomial regression model of the counts of cell types was used for testing for difference in cell type proportions between conditions, as described in Montoro et al., 2018. The negative binomial model was fitted using *glm.nb* from the MASS R package, and the *p*-value for the significance of the change in proportion between conditions was assessed using a likelihood-ratio test, computed using the *anova* function.

#### Gene expression plots

To compare gene expression in the fetal and adult datasets, raw data were merged using the concatenate function in Scanpy. Mean gene expression by cluster was displayed using the *dotplot* function in Scanpy. The percentage of cells within a cell type expressing a gene were indicated by dot size, and relative expression level was indicated by color intensity. Only dots with more than 10% of cells expressing the given gene were drawn.

#### Comparison between human and mouse DCs

Gene signatures were generated from publicly available datasets of murine splenic Xcr1^+^ DC (DC1) (*71*)(GSE71171), dermal migratory murine CD11b^+^ (DC2) (*74*)(GSE49358), human migratory tonsil and ascites DCs (*76*)(GSE115006 and GSE115007) as well as our own SmartSeq2 data from 3 rheumatoid arthritis synovial fluid donors. For each a list of DEGs (adjusted *p*-value <0.05) between the migratory DC cluster and all other MPs was generated to produce a gene signature. Enrichment scores of these signatures in adult skin was calculated using the *AddModuleScore* function in Seurat.

#### Cell-cell interaction using receptor-ligand analysis

CellPhoneDB v2.0 (*24*, *25*) was used to identify significant (*p* < 0.05) potential cell-cell ligand-receptor interactions for the entire healthy adult, AD and psoriasis datasets, using 1000 iterations of data permutations. A minimum threshold of expression proportion of 10% was set for any ligand or receptor in a given cell type.

#### Network analysis and clustered pathway annotation

We ranked the differential expression (log2fc) of genes between the analogous cell clusters in healthy and disease states using the Wilcoxon rank sum test with Benjamini and Hochberg method adjusted FDRs, for over-representation analysis (ORA) using G-profiler2 package in R to query two databases simultaneously (Reactome, Gene Ontology (GO) Biological Process) (**Table S5**). We derived statistical significance (*q* < 0.05) for each gene set enrichment and performed Markov clustering (MCL) using the MCL package in R to derive network neighborhoods based on shared genes between the gene sets. The gene set clusters were annotated using the AutoAnnotate Cytoscape package. Clusters were ranked by the mean enrichment score of all gene sets within each cluster and manually curated based on biological significance. We used the ClusterProfiler tool in R to visualize gene set enrichment.

#### Gene regulatory network prediction

We used the iRegulon package in Cytoscape to generate candidate transcription factors predicted to control homeostatic dendritic cell maturation. We ran the model using a search space of 20kb around the transcription start site and a ROC threshold for AUC calculation of 0.03.

#### Mass cytometry clustering analysis

Data from peeled, digested epidermis and dermis from 4 donors was clustered separately using the cytofkit Bioconductor package. Data was down-sampled to a maximum of 100,000 cells per experiment, clustered transformed using the cytofAsinh method, clustered with Rphenograph and visualized as tSNE. A manual gating strategy displayed in **Fig. S3** was used to overlay suggested cell identities onto the tSNE. Cell proportion bars were calculated as an average of 4 donors.

All p-values from statistical analyses are provided in **Table S6**.

---

**Fig. S1.**
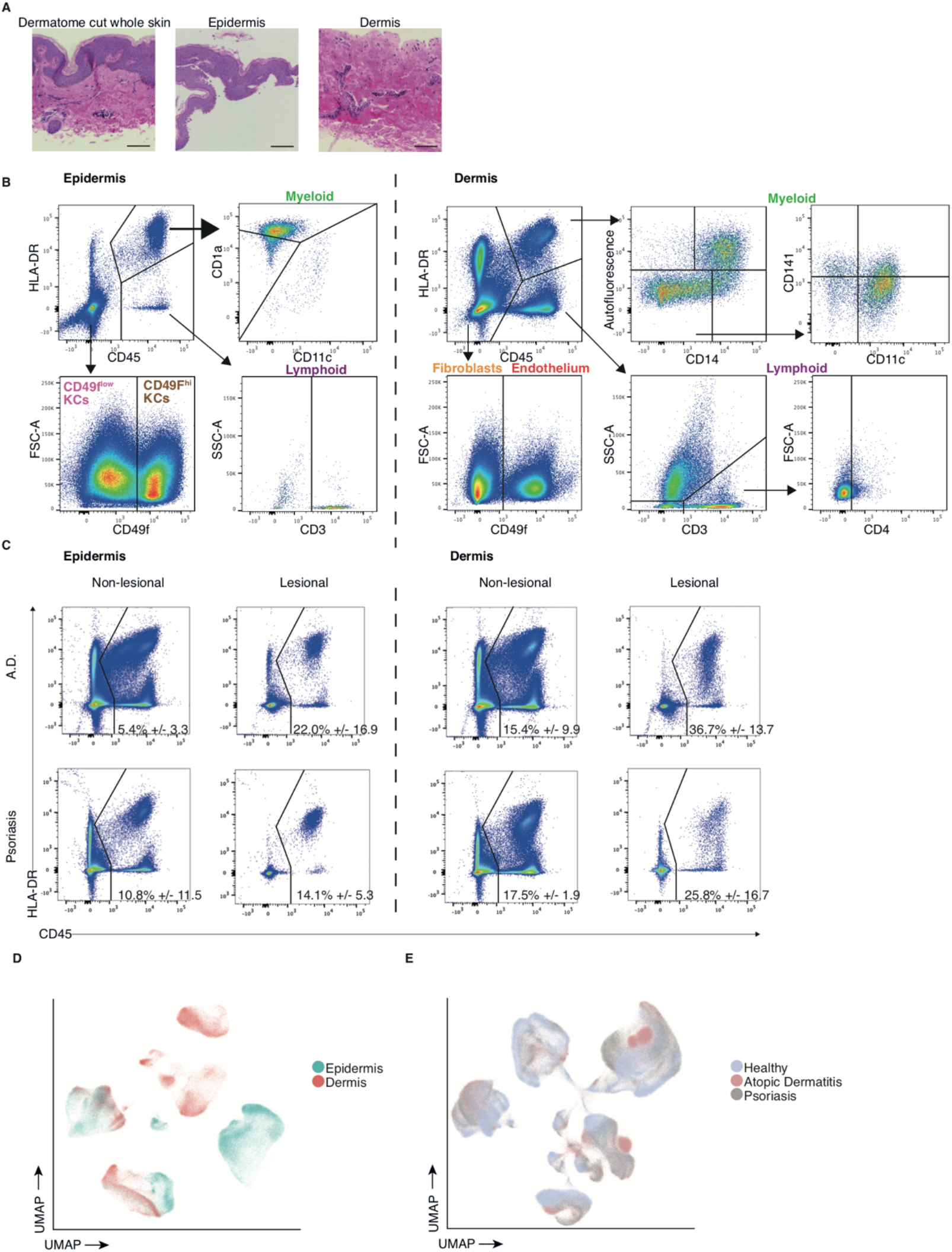
Skin dissociation and FACS enrichment. (a) Left: H&E stained image of skin fixed directly after dermatome cutting. Middle: Diagram of adult skin. Upper right: H&E stained image of epidermis fixed directly after peeling. Lower right: H&E stained image of dermis fixed directly after peeling. Scale bars represent 100μm. Images representative of n = 3. (b) FACS gating strategy used to collect cells for the healthy adult scRNA-seq experiments. Plots follow on from classical live singlet gating using DAPI and FSC-A/FSC-H respectively. (c) FACS gating strategy used to collect cells for the atopic dermatitis and psoriasis scRNA-seq experiments. Plots follow on from classical live singlet gating using DAPI and FSC-A/FSC-H respectively. (d) UMAP visualization shown in Figure 1B, colored by tissue origin of each cell. (e) UMAP visualization of all cells from adult healthy, atopic dermatitis and psoriasis samples, colored by disease state.

**Fig. S2.**
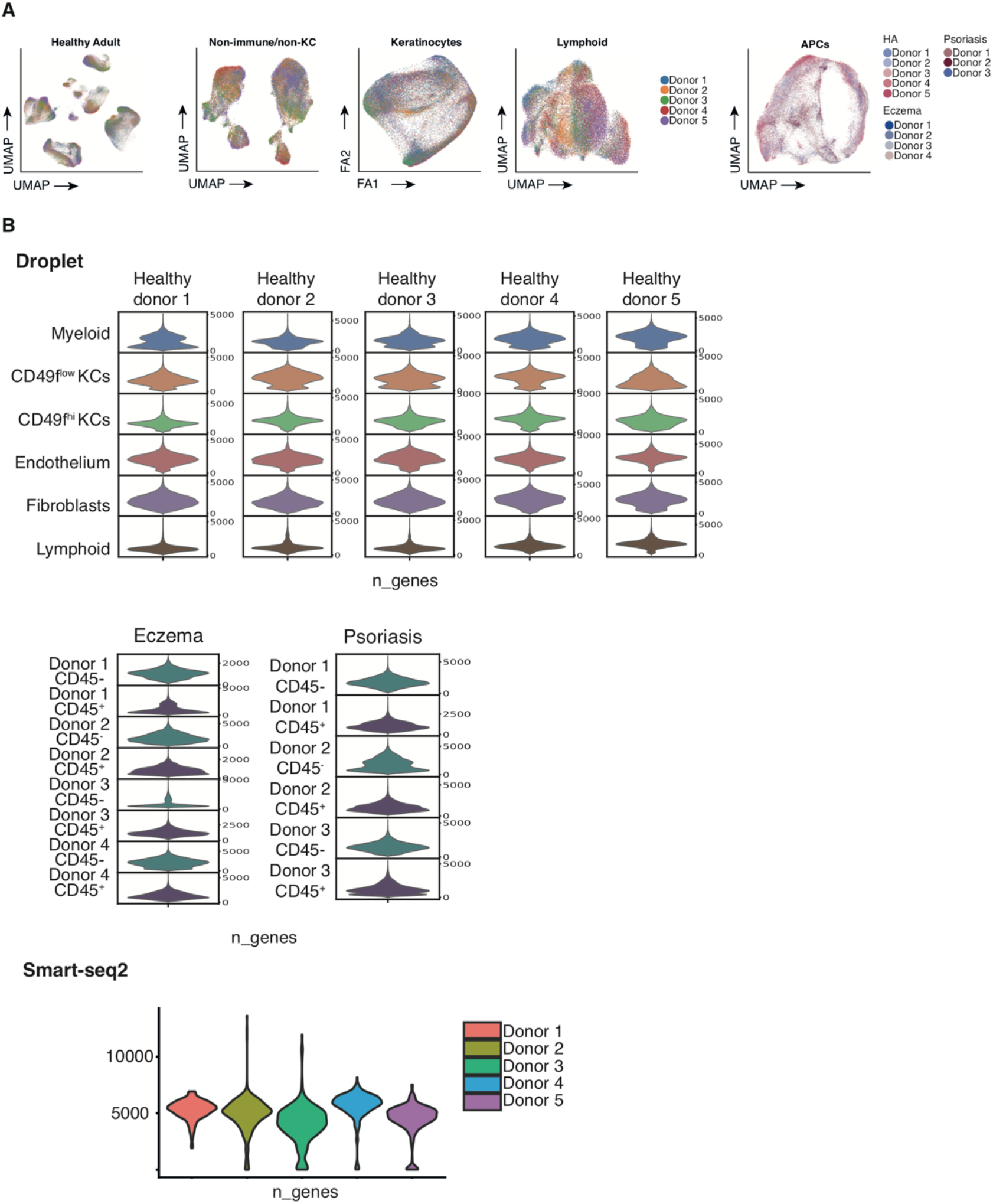
Single cell RNA seq quality control. (a) UMAP and force directed graph visualizations shown in each main figure, colored by sample origin of each cell. (b) Violin plots showing the number of UMIs detected in each sample for Droplet (10x Genomics) and Smart-seq2 scRNA-seq profiling.

**Fig. S3.**
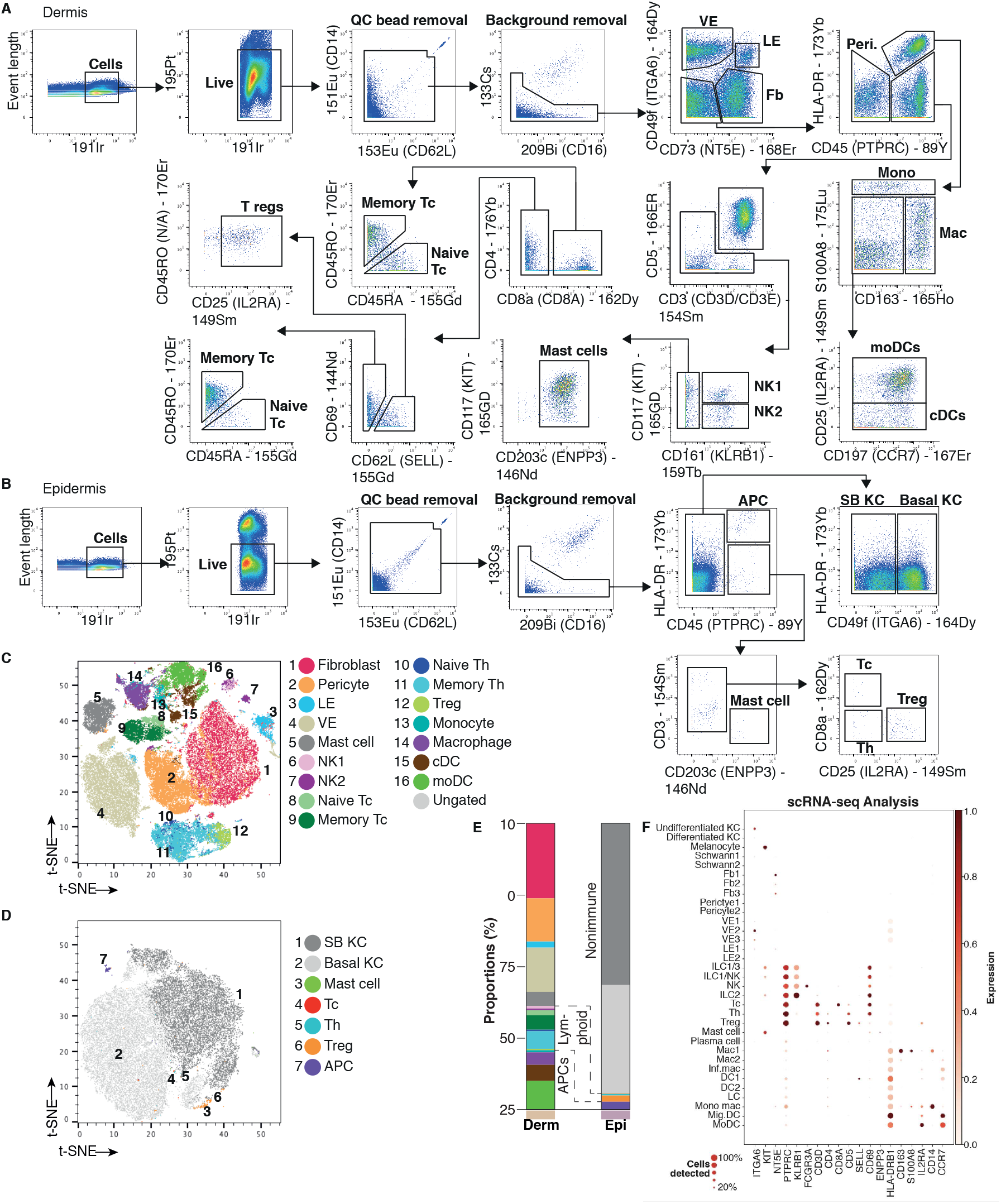
Suspension mass cytometry analysis of adult skin. (a) Gating strategy for dermal cells analyzed by mass cytometry. (b) Gating strategy for epidermal cells analyzed by mass cytometry. The same gating strategy from (a) overlaid onto a tSNE plot of 100,000 downsampled cells. (d) The same gating strategy from (b) overlaid onto a tSNE plot of 100,000 downsampled cells. (e) Bar chart showing the frequency of cells in each gate, cell type colors match legend in Supplementary Figure 3D. All plots representative of n=4. (f) Expression of marker genes that were used for mass cytometry panel in healthy adult skin cell states.

**Fig. S4.**
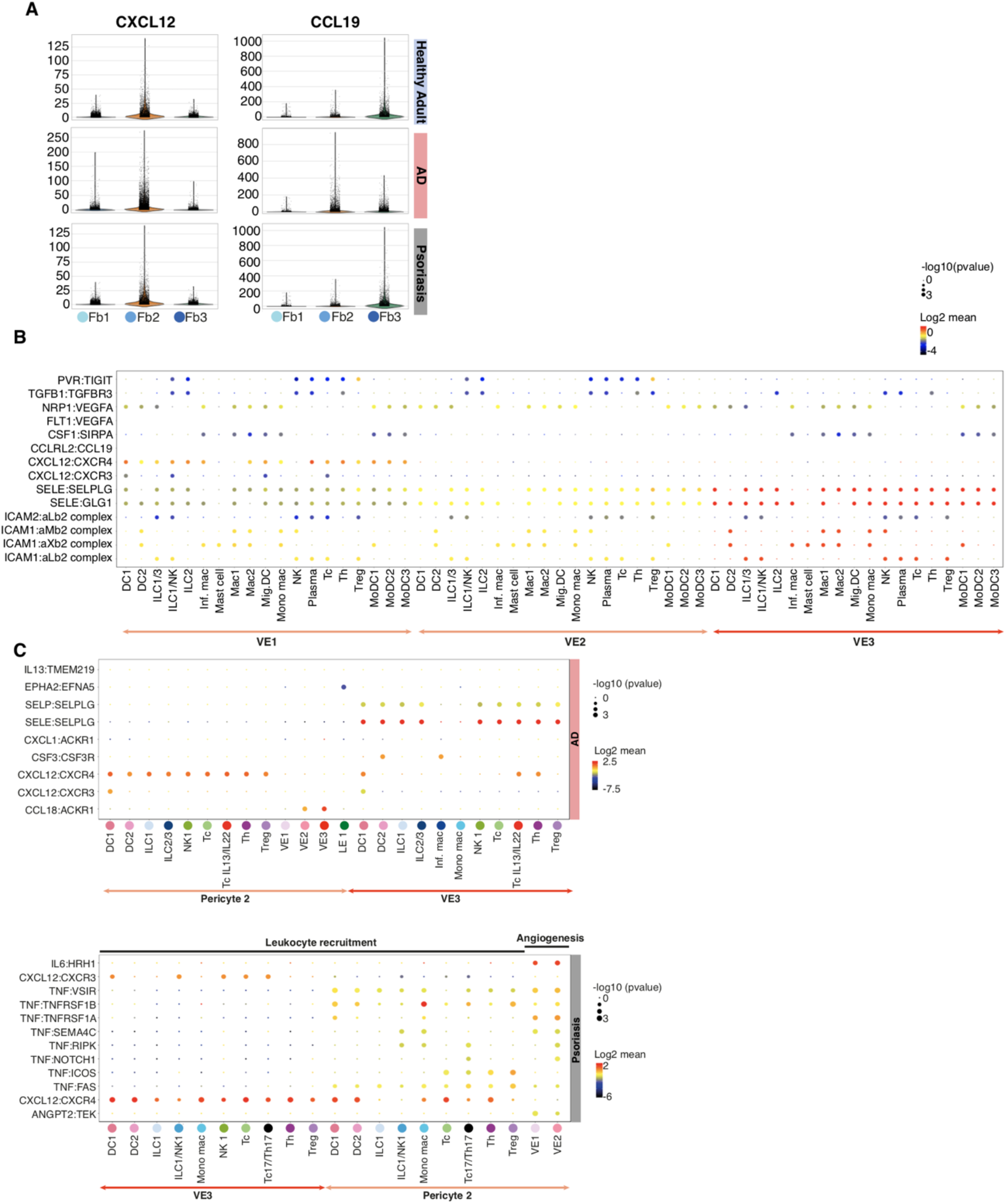
Stromal cells. (a) Violin plots of F1, F2, F3 expression for *CXCL12* and *CXCL19*. (b) CellPhoneDB predicted interactions between VE1, VE2 and VE3 and immune cells in healthy adult skin. (c) CellPhoneDB predicted interactions between VE3, pericyte2 and immune cells in AD (top panel) and psoriasis (bottom panel).

**Fig. S5.**
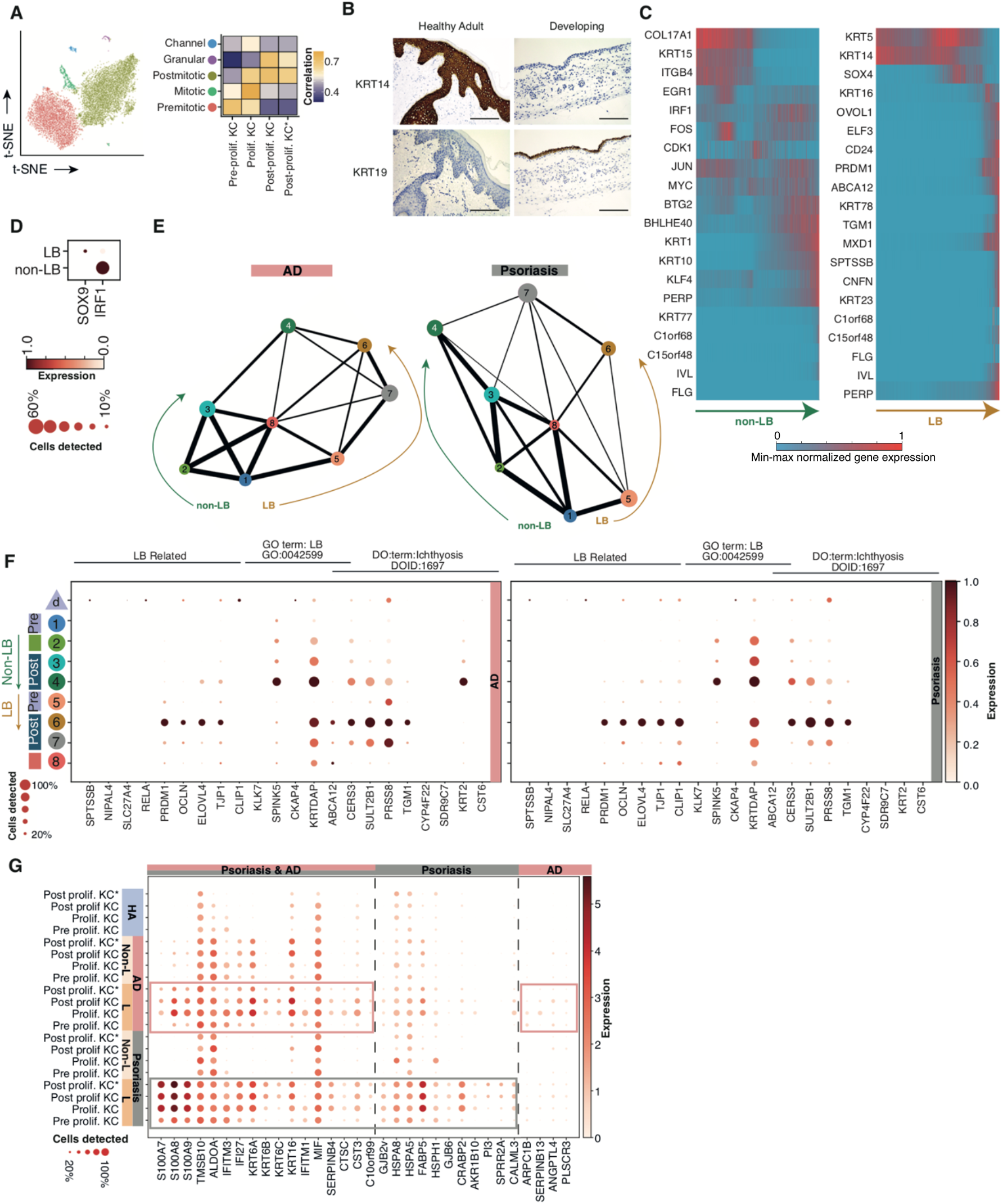
Keratinocytes. (a) Left panel: tSNE plot of keratinocytes isolated from published healthy breast reduction sample data (*8*), right panel: Spearman’s correlation between keratinocyte cell types of our data. (b) IHC staining of adult and fetal skin for keratin 14 and keratin 19. Scale bars represent 100μm. (c) Heatmaps of selected pseudotime genes from (**Fig. 3d**) across non-lamellar body expressing KCs (left) and lamellar body expressing KCs (right). (d) Dot plot showing the expression of *SOX9* and *IRF1* in the two basal keratinocyte differentiation pathways. (e) PAGA showing the relative connectivity between the keratinocyte clusters in AD (left panel) and psoriasis (right panel). Arrows indicate the two differentiation pathways of basal keratinocytes to suprabasal: LB = lamellar body. (f) Dot plot of genes related to lamellar body production and ichthyosis on the clusters presented in (**Fig. 3e**) as well as fetal keratinocytes. (g) Dotplot showing top differentially expressed genes for psoriasis (grey rectangle) and AD (pink rectangle) keratinocytes from lesional skin compared to healthy skin. HA = healthy adult (n=5), AD = atopic dermatitis (n=4), psoriasis (n=3), L = lesional, Non-L = non lesional.

**Fig. S6.**
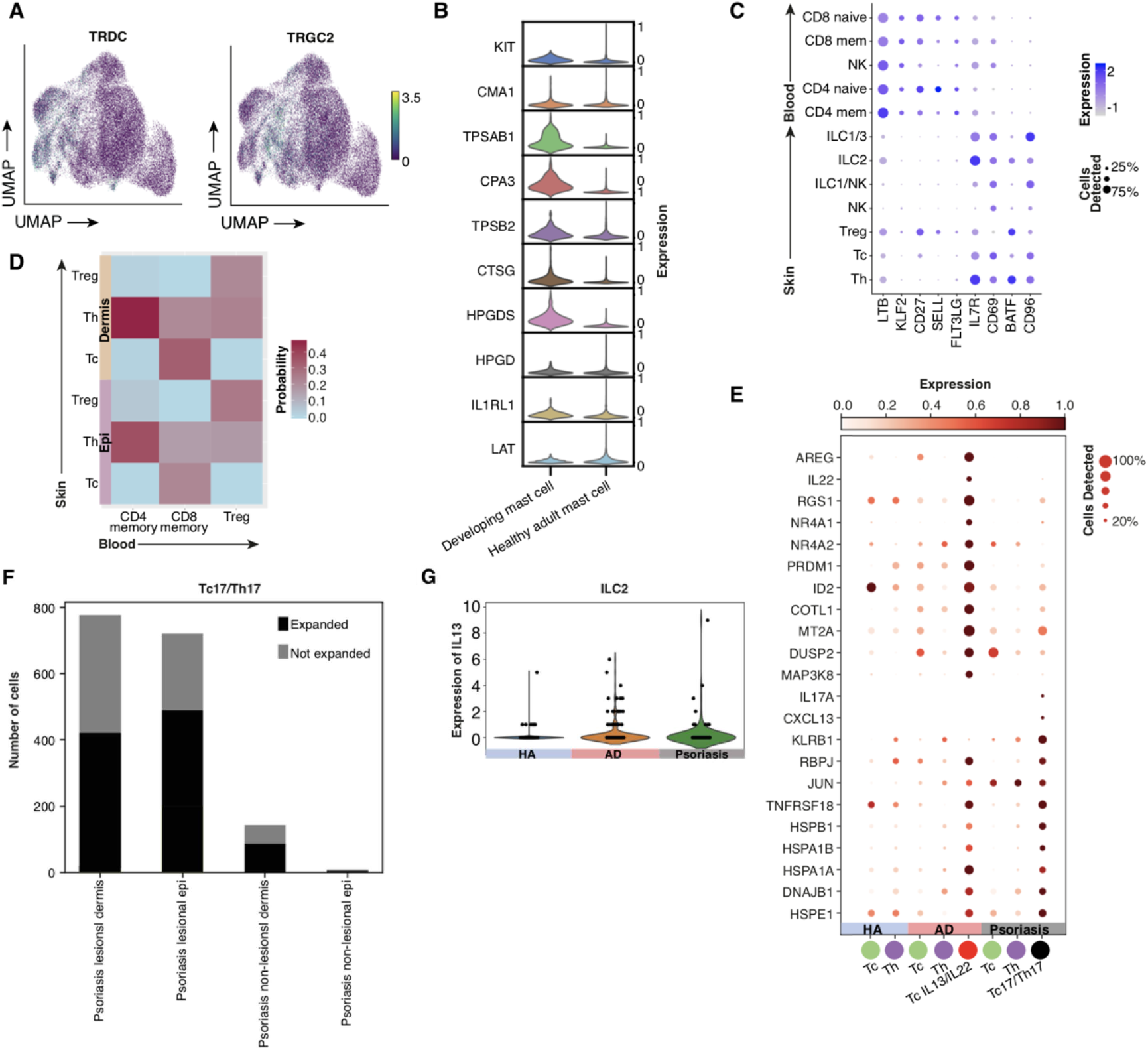
Lymphoid cells. (a) Feature plots showing γδ TCR gene expression on the UMAP plot shown in Figure 3A. (b) Violin plots showing the top 10 differentially expressed mast cell genes in fetal and adult skin mast cells. (c) Dot plot showing relative expression of skin residency related genes between blood and skin lymphocytes. (d) Heatmap of predicted closest match for blood T cells based on skin T cells as reference using the Transfer Anchors function in Seurat (e) Dot plot showing differentially expressed genes for T cell subtypes between healthy skin, AD and psoriasis. (f) Bar charts showing the number of expanded and not expanded Tc17/Th17 cells between psoriasis lesional and non lesional epidermis and dermis. P = psoriasis, L = lesional, NL = non lesional, Derm = dermis, Epi = epidermis. (g) Violin plot showing the expression of *IL13* in healthy adult, AD and psoriasis skin.

**Fig. S7.**
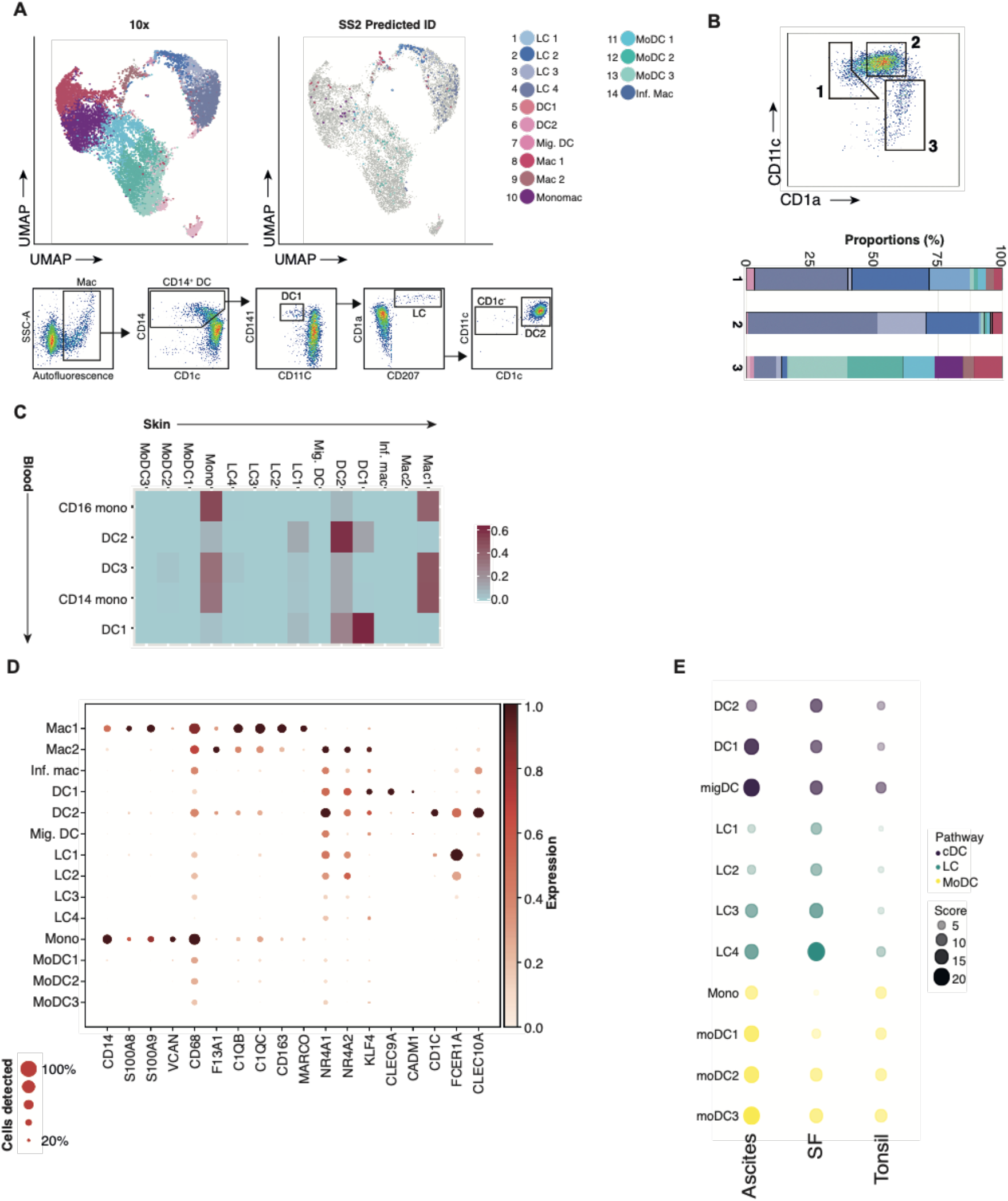
Mononuclear phagocytes I. (a) UMAP visualization of merged droplet and Smart-Seq2 data for skin MP. Annotation of Smart-seq2 data derived following projection on to droplet data using the Seurat TransferAnchors function. (b) Stacked bar graph of Smart-seq2 cells within each MP FACS-sort gate (c) Heatmap of predicted closest match for blood MPs based on skin MPs as reference using the Transfer Anchors function in Seurat (d) Dotplot of expression of specific marker genes for blood MPs by skin MPs (e) Dotplot of enrichment of gene signatures derived from migratory DCs from other human tissues.

**Fig. S8.**
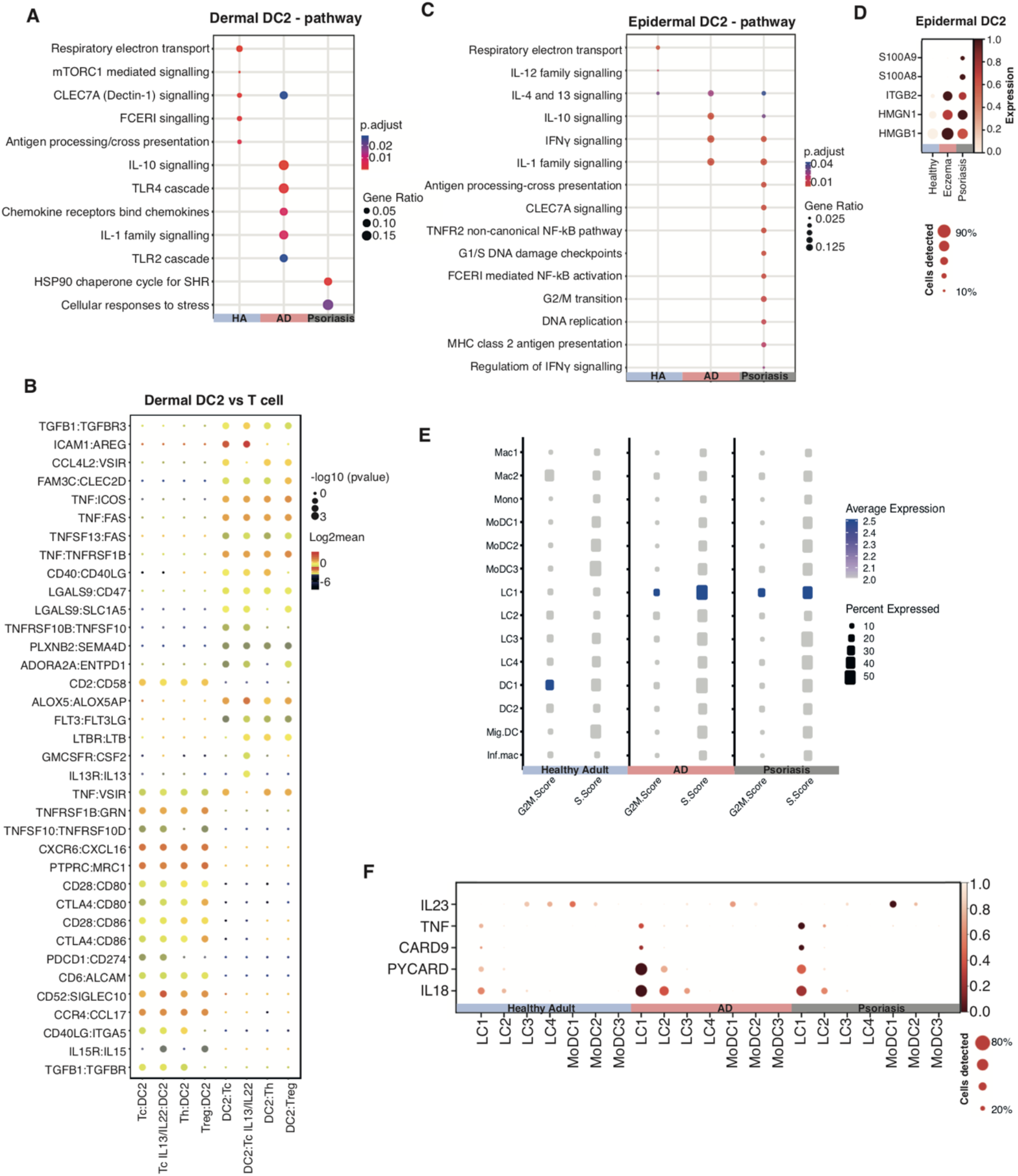
Mononuclear phagocytes II. (a) Dotplot of predicted functional modules within DEGs of dermal DC2 between health, AD and psoriasis. (b) CellPhoneDB predicted interactions between dermal DC2 and each

